# Amplification of potential thermogenetic mechanisms in cetacean brains

**DOI:** 10.1101/2020.10.23.352138

**Authors:** Paul R. Manger, Nina Patzke, Muhammad A. Spocter, Adhil Bhagwandin, Karl Æ. Karlsson, Mads F. Bertelsen, Abdulaziz N. Alagaili, Nigel C. Bennett, Osama B. Mohammed, Suzana Herculano-Houzel, Patrick R. Hof, Kjell Fuxe

## Abstract

To elucidate causality underlying the evolution of large brains in cetaceans, we examined the brains of 16 cetartiodactyl species for evidence of non-shivering thermogenesis. In comparison to the artiodactyl brain, the cetacean brain exhibits an expanded expression of uncoupling protein 1 (UCP1, UCPs being mitochondrial inner membrane proteins that dissipate the proton gradient to generate heat) in cortical neurons, localization of UCP4 within a substantial proportion of glia throughout the brain, and an increased density of noradrenergic axonal boutons (noradrenaline functioning to control concentrations of and activate UCPs). Thus, cetacean brains possess multiple characteristics indicative of intensified thermogenetic functionality that can be related to their current and historical obligatory aquatic niche. These findings necessitate reassessment of our concepts regarding the reasons for large brain evolution and associated functional capacities in cetaceans.

## Introduction

Cetaceans (whales, dolphins and porpoises) in general have large relative or absolute brain sizes, and are often considered cognitively complex mammals, their large brains apparently evolving in response to social and ecological demands present in their evolutionary history (Marino et al., 2008; Connor, 2007); however, alternative views regarding cetacean brain structure, function and evolution have been proposed (Kesarev, 1971; Nikolskaya, 2005; Manger, 2006, 2013; Patzke et al., 2015). The multiplicity of atypical features of the cetacean brain compared to other mammals (Kesarev, 1971; Manger, 2006; Patzke et al., 2015; Manger et al., 2010, 2012), their unusual sleep physiology (Lyamin et al., 2008), and that cetaceans have not been shown to outperform other mammals in behavioral tasks (Nikolskaya, 2005; Manger, 2013; Harley, 2013), have challenged the paradigm that cetaceans possess levels of cognitive complexity that differentiate them from the majority of other mammals.

It has been proposed that the current and historical, ubiquitous environmental pressure of water temperature has led to the evolution of the larger absolute or relative size of the cetacean brain (Manger, 2006). The mammalian brain is particularly sensitive to changes in temperature, with cortical neurons showing optimal functioning between 36-37°C, significantly decreased activity when brain temperature falls to 33°C, and loss of consciousness at 25-26°C (Mednikova et al., 2004). Thus, maintenance of brain temperature at levels appropriate for optimal neuronal functioning is an important aspect of mammalian physiology. Experimental evidence shows that exposure of the mammalian body to cold results in major decreases in body temperature but does not necessarily induce changes in brain temperature (Donhoffer, 1980). In addition, the temperature of the blood in the mammalian internal carotid artery is generally lower than that of the brain and jugular venous blood (Nybo et al., 2002; Vesterdorf et al., 2011). These studies indicate that the mammalian brain itself produces the heat required for optimal neuronal functioning, independent of thermogenetic mechanisms occurring in the remainder of the body. As there is no skeletal muscle within the mammalian cranial cavity, it is logical to posit that the production of heat by the brain would be through non-shivering thermogenetic mechanisms. Brown fat is a well-established site of non-shivering adaptive thermogenesis, and within brown fat, uncoupling proteins (UCPs) have been explicitly linked to the production of heat through their action on mitochondrial molecular pathways (Mao et al., 1999; Lowell and Spiegelman, 2000). Of the UCP family of proteins, all have been observed in the mammalian brain, but UCPs 1, 3, 4 and 5 are particularly strongly expressed and have been functionally linked to thermogenesis (Mao et al., 1999; Sanchis et al., 1998; Yu et al., 2000; Echtay, 2007). In addition, one of the many functions of noradrenaline is to control UCP concentrations and rapidly initiate UCP activity in brown adipocytes, leading to increased thermogenesis (Mory et al., 1984; Chunningham and Nicholls, 1987). Given the presence of UCPs and noradrenaline in the mammalian brain we examined the brains of three species of cetacean and eleven species of the closely related artiodactyls (even-toed ungulates) to explore the potential cellular basis of the thermogenetic hypothesis of cetacean brain evolution (Manger, 2006).

## Results

### Amplified UCP1 expression in cetaceans

Employing immunohistochemical techniques, UCP1 immunolocalization was observed in neocortical neurons in all cetartiodactyl species examined (Fig. 1, Table 1). Specificity of the UCP1 antibody was confirmed with Western blotting to brown fat taken from a laboratory rat (Fig. 2). UCP1 immunolabelling within the cortical neurons was observed in the perikaryal cytoplasm, as well as within the cytoplasm of the proximal portions of larger dendrites. The majority of the neurons immunopositive for UCP1 were pyramidal, although other cell types were also labelled (Fig. 1). Within artiodactyls, neurons immunopositive for UCP1 were observed mainly in the subgranular layers of the cerebral cortex (IV, V and VI) with occasional labelled neurons being observed in the supragranular cortical layers (I, II and III). In contrast, UCP1-immunopositive neurons were observed throughout all layers of the cetacean cerebral cortex. A systematic-random sampling analysis of the neurons immunopositive for UCP1 (Fig. 2, Table 1) revealed that the average percentage of neocortical neurons immunopositive for UCP1 in artiodactyls was 35.4% (range: 11.86% in blesbok anterior cingulate cortex to 58.25% in domestic pig anterior cingulate cortex, Table 1). In contrast, an average of 89.8% of cortical neurons were immunopositive for UCP1 in the cetacean cerebral cortex. The harbor porpoise (*Phocoena phocoena*) showed an average of 74.55% (range 71.28-83.19%) of cortical neurons being immunopositive for UCP1, while 100% of cortical neurons in the minke whale (*Balaenoptera acutorostrata*) and humpback whale (*Megaptera novaeangliae*) were immunopositive for UCP1 (Table 1). Using a two-proportions Z-test (as implemented in the R programming language) we tested the probability that the percentage of cortical neurons immunolabelled with UCP1 were equal in the artiodactyl and cetacean groups. Our analysis revealed that the proportion of immunolabelled UCP1 cortical neurons were significantly different between groups, with cetaceans having a significantly higher proportion of UCP1-immunoreactive neurons in both the occipital cortex (χ^2^ = 56.30; *P* = 6.21×10^−14^) and anterior cingulate cortex (χ^2^ = 51.69; *P* = 6.49×10^−13^) than the artiodactyls. These observations imply that there has been a proportional increase of UCP1 expression in the cortical neurons of cetaceans, to include almost all or all neurons of all layers, compared to artiodactyl where UCP1 expression is limited to a smaller proportion of neurons mostly within the subgranular cortical layers. In addition, UCP1-immunostained neurons were found throughout all grey matter regions of the harbor porpoise brain examined (Fig. 4).

**Table 1.** Specimens used and data generated in the current study. Brain masses, neuronal densities, grey and white matter glia densities, grey matter glia:neuron ratio, percentage (%) of grey matter/white matter neurons/glia immunopositive to uncoupling proteins 1 and 4 (UCP1, UCP4), density of boutons immunoreactive for dopamine-B-hydroxylase (DBH) and tyrosine hydroxylase (TH) in the grey matter and white matter, in anterior cingulate cortex (AC) and occipital cortex (OC).

| Species | Brain mass (g) | Neuronal density (mm<sup>3</sup>) |  | Grey matter glia density (mm<sup>3</sup>) |  | Glia:neuron ratio |  | White matter glia density (mm<sup>3</sup>) |  | % grey matter neurons UCP1 immunopositive |  | % grey matter glia UCP4 immunopositive |  | % white matter glia UCP4 immunoreactive |  | Grey matter DBH bouton density (mm<sup>3</sup>) |  | White matter DBH bouton density (mm<sup>3</sup>) |  | Grey matter TH bouton density (mm<sup>3</sup>) |  | White matter TH bouton density (mm<sup>3</sup>) |  |
| --- | --- | --- | --- | --- | --- | --- | --- | --- | --- | --- | --- | --- | --- | --- | --- | --- | --- | --- | --- | --- | --- | --- | --- |
|  |  | AC | OC | AC | OC | AC | OC | AC | OC | AC | OC | AC | OC | AC | OC | AC | OC | AC | OC | AC | OC | AC | OC |
| Artiodactyls |  |  |  |  |  |  |  |  |  |  |  |  |  |  |  |  |  |  |  |  |  |  |  |
| *Gazella marica* | 63.3 | 13967 | 22184 | 112892 | 220405 | 6.820 | 6.030 | - | - | 58.19 | 36.28 | 0 | 0 | 0 | 0 | 8218 | 11077 | 1325 | 1900 | 6132 | 9405 | 2538 | 1750 |
| *Sus scrofa* | 64.0 | 18355 | 23808 | 108238 | 133283 | 5.897 | 5.598 | 273326 | 230191 | 58.25 | 57.69 | 0 | 0 | 0 | 0 | 9257 | 10682 | 2900 | 2988 | 5618 | 8595 | 1675 | 1013 |
| *Capra nubiana* | 132.4 | 12992 | 13104 | 81151 | 87317 | 6.246 | 6.663 | - | - | 27.72 | 29.00 | 0 | 0 | 0 | 0 | 6658 | 7229 | 2200 | 3025 | 6560 | 7561 | 1200 | 913 |
| *Antidorcas marsupialis* | 224.2 | 11808 | 13887 | 69440 | 86121 | 5.881 | 6.202 | - | - | 23.12 | 38.21 | 0 | 0 | 0 | 0 | 7699 | 7000 | 2350 | 2300 | 6347 | 7717 | 1675 | 1025 |
| *Damaliscus pygargus* | 230.0 | 19632 | 28189 | 83679 | 105486 | 3.740 | 4.260 | 283841 | 250063 | 11.86 | 30.35 | 0 | 0 | 0 | 0 | 7258 | 7856 | 2288 | 1288 | 4073 | 8135 | 1450 | 1450 |
| *Tragelaphus strepsiceros* | 355.0 | 13771 | 14849 | 52950 | 71238 | 4.790 | 3.840 | 205760 | 180886 | 21.16 | 45.10 | 0 | 0 | 0 | 0 | 6595 | 10965 | 1688 | 2683 | 5077 | 10097 | 2650 | 838 |
| *Connochaetes taurinus* | 385.0 | 22108 | 22957 | 70145 | 76245 | 3.173 | 3.321 | 221160 | 176321 | 53.84 | 24.27 | 0 | 0 | 0 | 0 | 6670 | 11390 | 2188 | 2763 | 4763 | 9313 | 1363 | 675 |
| *Camelus dromedarius* | 395.0 | 15635 | 9930 | 99084 | 66354 | 6.337 | 6.682 | - | - | 40.91 | 32.64 | 0 | 0 | 0 | 0 | 6478 | 10887 | 2675 | 3600 | 5785 | 10815 | 2000 | 538 |
| *Tragelaphus angasii* | 417.2 | 9432 | 10718 | 65040 | 87719 | 6.896 | 8.184 | - | - | 35.13 | 41.60 | 0 | 0 | 0 | 0 | 10278 | 11233 | 2875 | 3788 | 7677 | 10770 | 1950 | 1425 |
| *Hippopotamus amphibius* | 435.5 | 7519 | 8135 | 58984 | 75775 | 7.848 | 9.315 | - | - | 23.26 | 33.71 | 0 | 0 | 0 | 0 | 9867 | 12305 | 3400 | 2488 | 8320 | 11704 | 3288 | 2225 |
| *Syncerus caffer* | 514.8 | 13889 | 16739 | 106311 | 111595 | 7.654 | 6.667 | - | - | 20.61 | 36.28 | 0 | 0 | 0 | 0 | 8970 | 12900 | 2513 | 2113 | 8172 | 8483 | 1400 | 863 |
| Cetaceans |  |  |  |  |  |  |  |  |  |  |  |  |  |  |  |  |  |  |  |  |  |  |  |
| *Phocoena phocoena* | 486.0 | 10938 | 20129 | 97623 | 153415 | 7.620 | 8.930 | 210627 | 187064 | 78.59 | 63.96 | 33.27 | 31.84 | 63.45 | 47.17 | 11763 | 16013 | 2475 | 1613 | 13973 | 18247 | 1875 | 2200 |
| *Phocoena phocoena* | 502.0 | 10938 | 20129 | 97623 | 153415 | 8.530 | 7.980 | 202072 | 239250 | 83.19 | 72.45 | 32.74 | 33.58 | 69.17 | 45.06 | 11567 | 15898 | 2825 | 2063 | 14092 | 17802 | 1625 | 2088 |
| *Balaenoptera acutorostrata* | 2600.0 | 8671 | 11119 | 82149 | 114384 | 9.470 | 10.290 | 160172 | 175982 | 100.00 | 100.00 | 58.88 | 39.77 | 50.72 | 64.73 | 9156 | 13488 | 4163 | 2138 | 11786 | 15010 | 1613 | 1738 |
| *Balaenoptera acutorostrata* | 3200.0 | 8671 | 11119 | 82149 | 114384 | 12.160 | 8.280 | 182841 | 231483 | 100.00 | 100.00 | 35.24 | 31.88 | 50.31 | 54.45 | 10208 | 13303 | 1513 | 4963 | 10717 | 14846 | 1875 | 1750 |
| *Megaptera novaeangliae* | 4600.0 | - | 6922 | - | 183231 | - | 10.220 | - | 183231 | 100.00 | 100.00 | - | 29.33 | - | 58.97 | - | - | - | - | - | - | - | - |

**Fig. 1.**
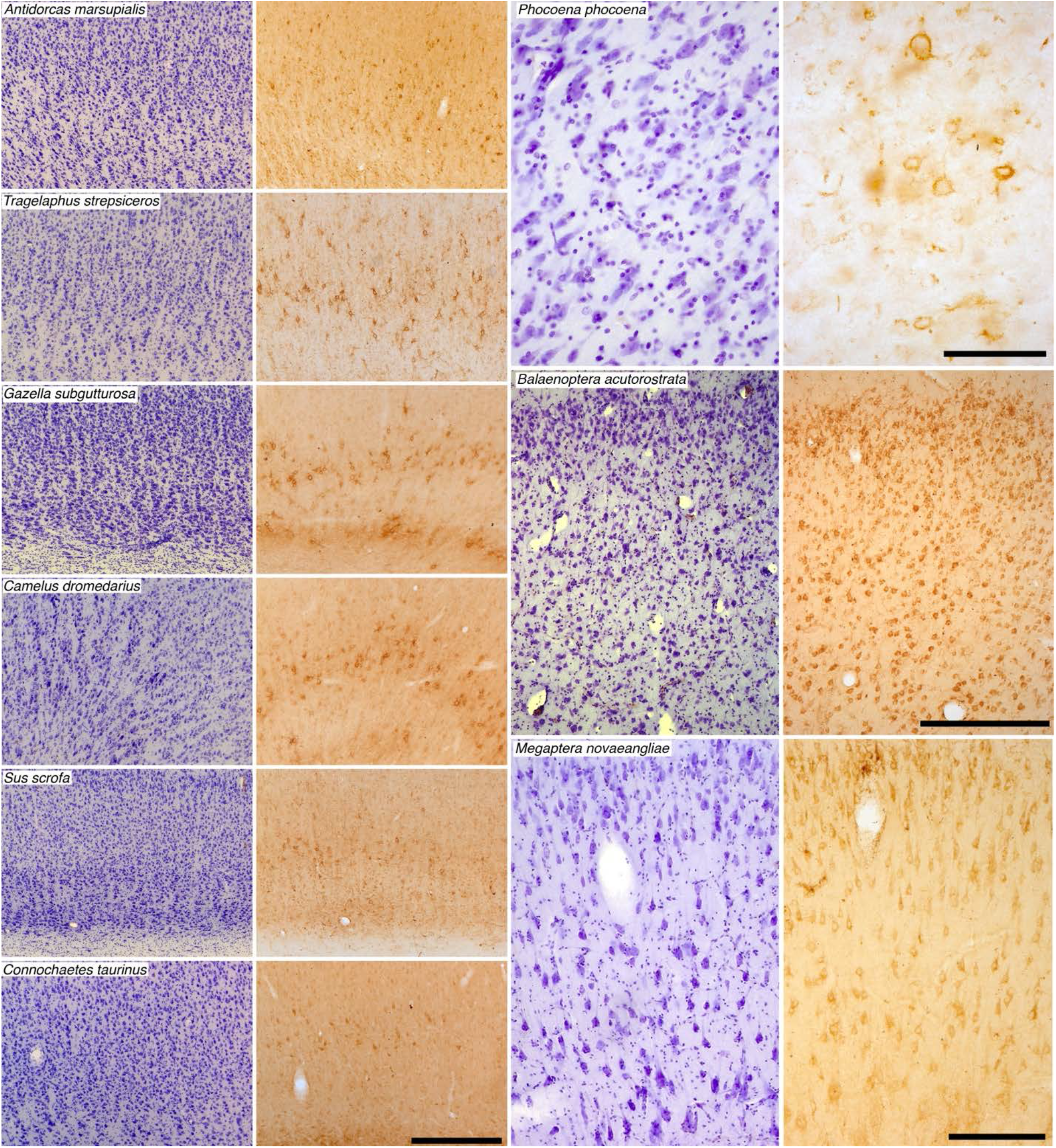
UCP1 immunostaining in cetartiodactyl cerebral cortex. Photomicrographs of Nissl stained (purple colored images) and UCP1 immunostained (brown colored images) cortical sections in a range of artiodactyl (two left columns) and cetacean species (two right columns). Note in all cases the presence of UCP1 immunostained cortical neurons, but in the artiodactyls these are limited to the lower layers of the cortex, while almost all cortical neurons from all layers are immunopositive in the cetaceans. Scale bar in the UCP1 stained section of *Connochaetes taurinus* equals 500 μm and applies to all artiodactyl images. Scale bar in the UCP1 stained section of *Phocoena phocoena* equals 100 μm and applies to both images. Scale bar in the UCP1 stained section of *Balaenoptera acutorostrata* equals 500 μm and applies to both images. Scale bar in the UCP1-immunostained stained section of *Megaptera novaeangliae* equals 250 μm and applies to both images.

**Fig 2.**
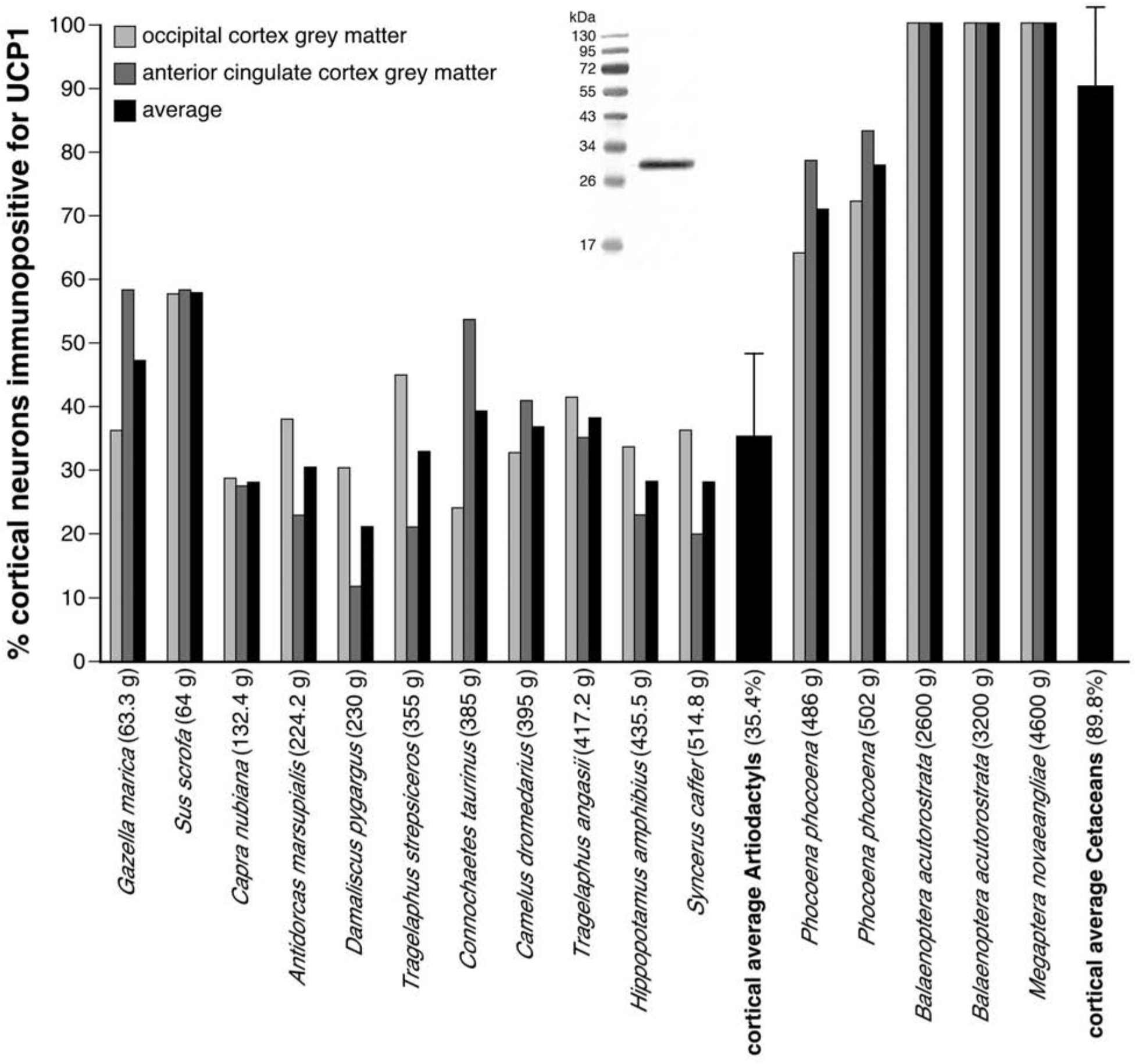
Quantification of UCP1 immunostaining in cetartiodactyl cerebral cortex. Graphical representation of the results of the stereological analysis of the percentage of cortical neurons immunopositive for UCP1 in the occipital and anterior cingulate cortices of the species studied. For each species the brain mass is given in grams next to the name on the x-axis. Note that the average percentage of cortical neurons immunopositive for UCP1 in the artiodactyls studied was 35.4%, while in the cetaceans studied it was 89.9% (Table 1, error bars on average bars represent one standard deviation). The Western immunoblot in the middle of the graph shows the specificity of the UCP1 antibody to brown fat taken from a laboratory rat.

### UCP4/5 expression in cetacean glia

UCP4 has been identified using Northern (RNA) blots in the human brain and is suggested to play a role in thermogenesis (Mao et al., 1999). Using Western blots, we found evidence for the presence of UCP4 in the brains of all artiodactyl and cetacean species studied (Fig. 3). In contrast to the detectable presence of UCP4 with Western blotting, immunohistochemical localization of UCP4 was only observed in the cetacean brains. In all three cetacean species studied, we observed strong immunolocalization of UCP4, and weaker immunolocalization of UCP5, within glial cells in the cerebral cortex and the subcortical white matter, but no staining of neurons (Fig. 3, Table 1). In the harbor porpoise an average of 33.16% of glial cells in the cerebral cortical grey matter (from anterior cingulate and occipital regions) were immunopositive for UCP4, while an average of 57.12% of glial cells in the cortical white matter (from anterior cingulate and occipital regions) were immunopositive for UCP4. In the minke whale an average of 41.44% of glial cells in the cortical grey matter and an average of 55.05% of glial cells in the cortical white matter were immunopositive for UCP4. In the humpback whale an average of 29.33% of glial cells in the cortical grey matter and an average of 58.97% of glial cells in the cortical white matter were immunopositive for UCP4. Thus, in cetaceans, approximately 36% of glial cells in the cortical grey matter and 56% of glial cells in the cortical white matter show specific immunolocalization of UCP4 (Table 1). In all three species UCP5 was also expressed in similar proportions of glial cells, but the strength of immunostaining was substantially weaker. A limited examination of the immunolocalization of UCP4 and UCP5 in other regions of the harbor porpoise brain showed similar levels of glial staining in both grey and white matter (Fig. 4), indicating that UCP4 and UCP5 are proteins likely to be expressed in glial cells throughout the entire cetacean brain. Based on these observations, we conclude that while UCP4 and UCP5 are proteins found in the brains of both artiodactyl and cetacean species, in cetaceans they exhibit a specific localization to glial cells, indicating a specialization in their expression, and related function, in cetaceans.

**Fig. 3.**
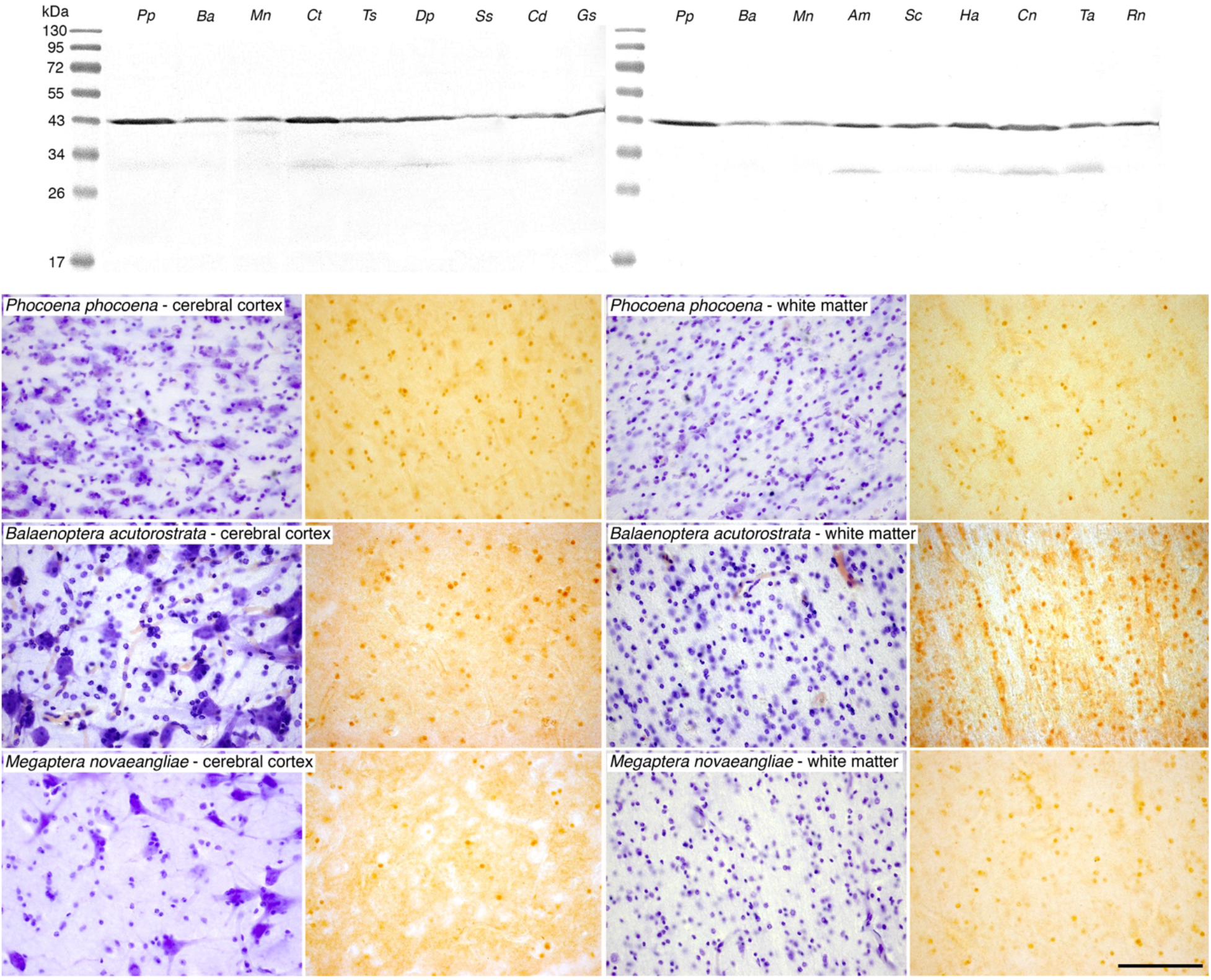
UCP4 Western blotting and immunostaining in cetartiodactyl cerebral cortex. While UCP4 was present in the cortical grey and white matter of all species, as evidenced in the Western blot at the top of the panel, it was only found to be immunolocalized to glial cells in the cetaceans. Photomicrographs of Nissl-stained (purple colored images) and UCP4-immunostained (brown colored images) from cortical and subcortical white matter sections in a range of cetacean species. Note the presence of UCP4-immunoreactivity in approximately 30% of glial cells in the cerebral cortex and approximately 60% of glial cells in the white matter in all cetacean species (Table 1). The scale bar in the UCP4 stained section of *Megaptera novaeangliae* – white matter, equals 100 μm and applies to all photomicrographs. ***Pp*** – harbor porpoise, *Phocoena phocoena*; ***Ba*** – minke whale, *Balaenoptera acutorostrata*; ***Mn*** – humpback whale, *Megaptera novaeangliae*; ***Ct*** – blue wildebeest, *Connochaetes taurinus*; ***Ts*** – greater kudu, *Tragelaphus strepsiceros*; ***Dp*** – blesbok, *Damaliscus pygargus*; ***Ss*** – domestic pig, *Sus scrofa*; ***Cd*** – dromedary camel, *Camelus dromedarius*; ***Gm*** – sand gazelle, *Gazella marica*; ***Am*** – springbok, *Antidorcas marsupialis*; ***Sc*** – African buffalo, *Syncerus caffer*; ***Ha*** – river hippopotamus, *Hippopotamus amphibius*; ***Cn*** – Nubian ibex, *Capra nubiana*; ***Ta*** – nyala, *Tragelaphus angasii*; ***Rn*** – laboratory rat, *Rattus norvegicus*.

**Fig. 4.**
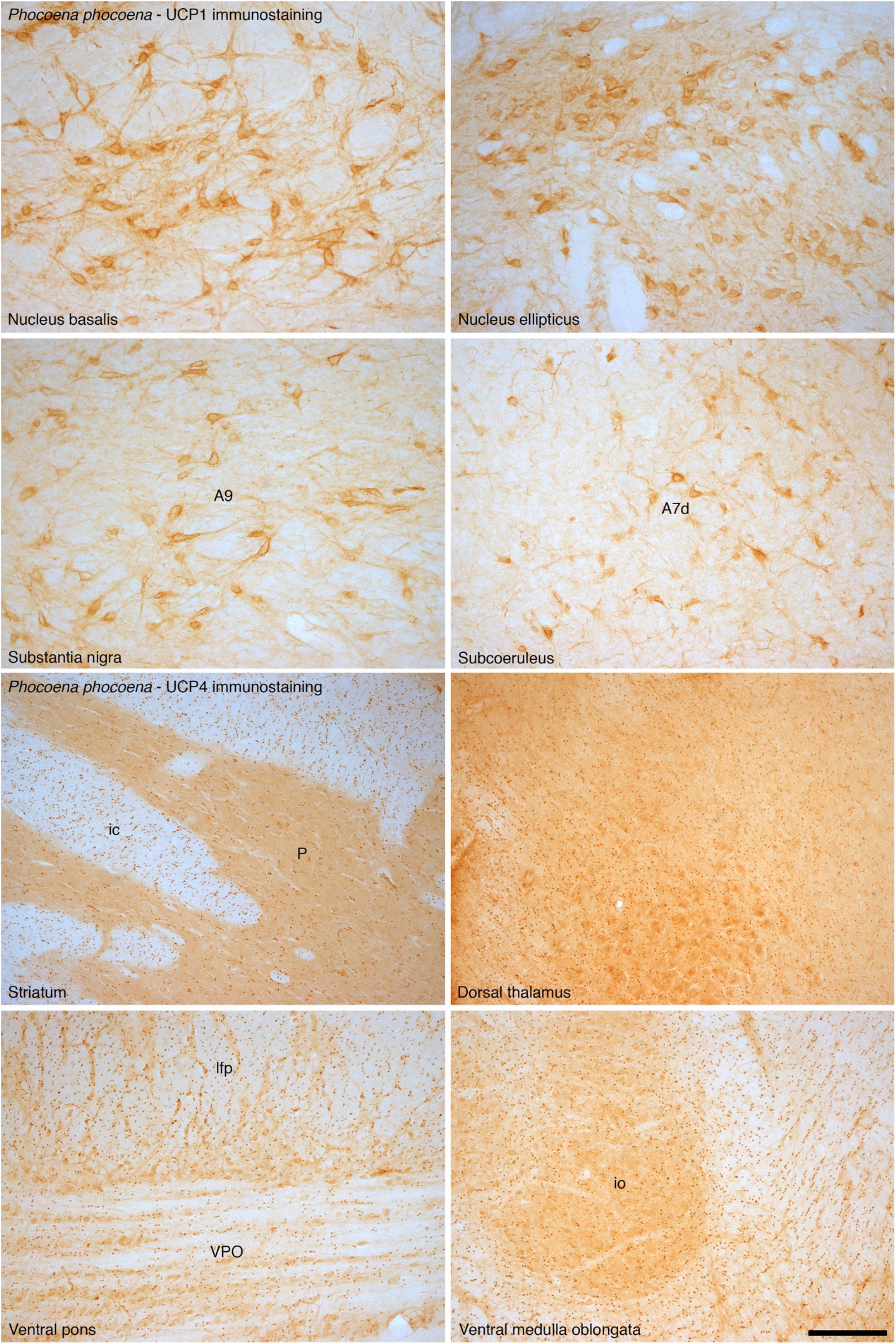
UCP1 and UCP4 immunostaining in non-cortical regions of the harbor porpoise brain. In addition to examining the expression of UCP1 and UCP4 in the cerebral cortex of the brain of the harbor porpoise, we examined several other brain regions. In all regions we found neurons with distinct UCP1 immunoreactivity, with an intracellular staining pattern similar to that observed in the neurons of the cerebral cortex. The photomicrographs shown here depict UCP1 immunostaining in various non-cortical regions of the harbor porpoise brain, including the nucleus basalis, nucleus ellipticus, the substantia nigra (**A9**), and the nucleus subcoeruleus (**A7d**, its diffuse region). In addition, in all regions we found glial cells with distinct immunoreactivity to the UCP4 antibody. Interestingly, the density of glial cells immunopositive for UCP4 appears higher in the white matter than in the grey matter, reflecting the same proportional distribution of stained glia as when comparing the white and grey matter of the cerebral cortex. The photomicrographs shown here depict UCP4 immunostaining in various non-cortical regions of the harbor porpoise brain, including the striatum (**P** – putamen, **ic** – internal capsule), dorsal thalamus, ventral pons (**VPO** – ventral pontine nucleus, **lfp** – longitudinal fasciculus of pons) and the ventral medulla oblongata (**io** – portion of inferior olivary nuclear complex). Scale bar = 250 μm and applies to all.

### Noradrenergic bouton density in cetacean cerebral cortex

As one of the many known functions of noradrenaline (NA) is to control concentrations of UCPs and initiate UCP activity in brown adipocytes (Mory et al., 1984; Cunningham and Nicholls, 1987), we used immunohistochemical staining for dopamine-β-hydroxylase (DBH, the enzyme that converts dopamine to noradrenaline in the catecholamine biosynthetic pathway) to examine the density of noradrenergic boutons in the grey and white matter of the cerebral cortex (from anterior cingulate and occipital regions) in the cetartiodactyl species studied (Fig. 5; Table 1). The average density of NA boutons in the cortical grey matter of the artiodactyls studied was 8980 boutons/mm^3^ (range: 6478/mm^3^ in dromedary camel anterior cingulate cortex to 12900/mm^3^ in African buffalo occipital cortex). In cetacean cortical grey matter, an average density of 12675 NA boutons/mm^3^ was observed (range: 9156/mm^3^ in minke whale anterior cingulate cortex to 16013/mm^3^ in harbor porpoise occipital cortex, Table 1). Using a two sample T-test we compared DBH-immunoreactive bouton density in the grey matter of the anterior cingulate and occipital cortex between artiodactyls and cetaceans. Cetaceans have significantly higher DBH-immunoreactive bouton densities in both the anterior cingulate and occipital cortex compared to artiodactyls (anterior cingulate: *t* = −3.595; df =15, *P* = 0.011; occipital cortex: *t* = - 4.546; df =15, *P* = 0.002). In the cortical white matter, an average density of 2515 NA boutons/mm^3^ was observed in artiodactyls, which was not significantly different to (anterior cingulate: *t* =-0.5977; df =15, *P* = 0.585; occipital: *t* = −0.08; df =15, *P* = 0.941) the average NA bouton density found in cetacean cortical white matter (2719 NA boutons/mm^3^, Table 1, Fig. S1). When a third variable, such as cortical neuron density, cortical glia density or brain mass (Table 1) were analyzed with the current data using analysis of covariance (ANCOVA), cetaceans were still observed to have statistically significantly higher DBH-immunoreactive bouton densities in the cortical grey matter than artiodactyls. Thus, in addition to having an amplified (UCP1) and localized (UCP4/5) representation of UCPs in the cortical grey matter, the cetaceans have a significantly denser noradrenergic innervation, which likely functions to increase concentrations of, and activate, UCPs. Quantitative analysis of bouton densities following immunohistochemical staining for tyrosine hydroxylase (TH, the enzyme that converts tyrosine to L-3,4-dihydroxyphenylalanine in the catecholamine biosynthetic pathway) provided similar results (Figs. S2, S3, S4).

**Fig. 5.**
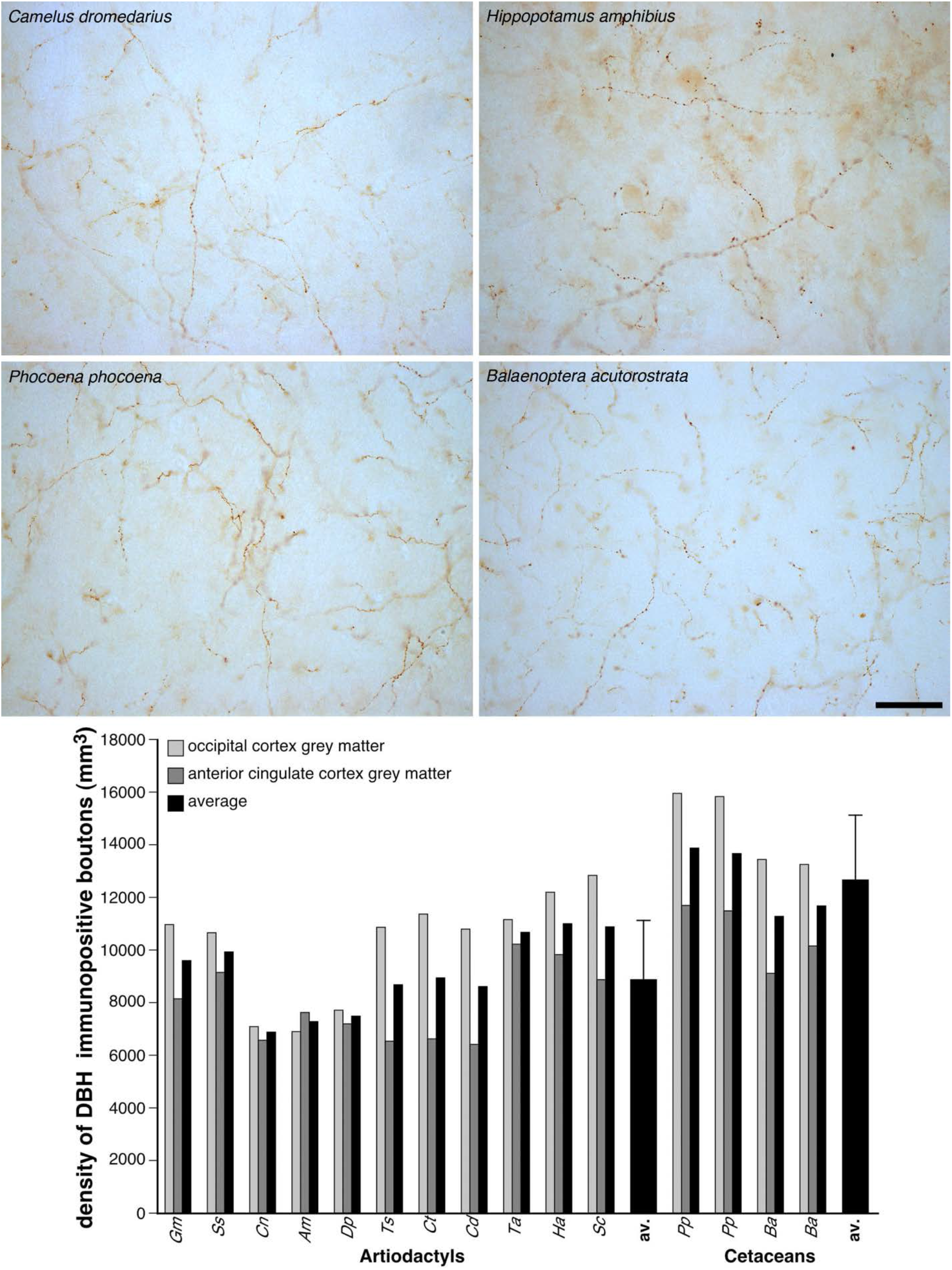
Quantification of noradrenergic bouton density in cetartiodactyl cerebral cortex. Photomicrographs of dopamine-ß-hydroxylase (DBH)-immunostained axonal boutons in the occipital cortical grey matter of *Camelus dromedarius*, *Hippopotamus amphibius*, and *Phocoena phocoena*, and the anterior cingulate cortical grey matter of *Balaenoptera acutorostrata*. The scale bar = 50 μm and applies to all photomicrographs. Note the higher density of the DBH-immunoreactive boutons in the cortical grey matter of cetaceans compared to the artiodactyls as confirmed with stereological analysis (see the graph below the photomicrographs), showing that the density of DBH-immunoreactive boutons in the cortical grey matter of cetaceans is, on average, 1.4 times higher than that observed in artiodactyls (Table 1, error bars on average bars represent one standard deviation). ***Gm*** – sand gazelle, *Gazella marica*; ***Ss*** – domestic pig, *Sus scrofa*; ***Cn*** – Nubian ibex, *Capra nubiana*; ***Am*** – springbok, *Antidorcas marsupialis*; ***Dp*** – blesbok, *Damaliscus pygargus*; ***Ts*** – greater kudu, *Tragelaphus strepsiceros*; ***Ct*** – blue wildebeest, *Connochaetes taurinus*; ***Cd*** – dromedary camel, *Camelus dromedarius*; ***Ta*** – nyala, *Tragelaphus angasii*; ***Ha*** – river hippopotamus, *Hippopotamus amphibius*; ***Sc*** – African buffalo, *Syncerus caffer*; **av.** – average; ***Pp*** – harbor porpoise, *Phocoena phocoena*; ***Ba*** – minke whale, *Balaenoptera acutorostrata*.

## Discussion

### Augmented thermogenic features of cetacean brains

Our observations indicate that the cetacean brain appears to house three augmented characteristics of a pre-existing, brain-based, non-shivering thermogenic system that should increase heat generation capabilities above and beyond that seen in the brains of other mammals. First, the expanded expression of UCP1 throughout almost all cortical neurons indicates that, unlike in artiodactyls, the majority of cortical neurons within the cetacean brain can function as thermogenic units if necessary. Second, the specific localization of UCP4/5, within many glial cells of the cetacean brain, indicates that between 30 and 70% of glial cells may be employed as thermogenic units in both grey and white matter if necessary. The generally higher density of glial cells and the higher glia:neuron ratio in the cetacean brain (Table 1) indicates that glial based UCPs may form a potentially powerful thermogenic mechanism in the cetacean brain. Last, the increased density of noradrenergic boutons in the cetacean cerebral cortex compared to the artiodactyls indicates that the capacity to increase concentrations of UCP within the tissue and activate these proteins appears to be enhanced in the cetaceans compared to the artiodactyls. As cetaceans undergo unihemispheric slow wave sleep (USWS) without rapid eye movement sleep (Lymain et al., 2008), and have a very higher number of noradrenergic neurons in the locus coeruleus complex (Dell et al., 2016a,b) (Fig. S5), the cetacean brain is likely to have a steady supply of noradrenaline, which will not occur in the artiodactyls, again enhancing the potential for thermogenesis by the cetacean brain. This link between noradrenaline and thermogenesis in the cetacean brain is supported by the observation that during cetacean USWS, when the activity of the noradrenergic neurons of the locus coeruleus complex (Fig. S5) is reduced unilaterally (Ridgway et al., 2006), the temperature of the ipsilateral sleeping hemisphere gradually decreases (Lyamin et al., 2008). In summary, neuronal and glial portions of the cetacean brain appear to have specializations associated with UCPs, and the increased noradrenergic input should act to increase the concentration of these proteins in the tissue and activate them. This indicates that amplified thermogenetic capabilities are likely to be an extremely important basic function of the cetacean central nervous system (Manger, 2006).

The findings presented herein support the thermogenesis hypothesis of cetacean brain evolution and function (Manger, 2006, 2013). The presence of UCPs in the majority of cortical neurons as well as within a substantial proportion of glial cells, together with the associated increased of noradrenergic innervation throughout the grey and white matter of the brain is important, because in situations of thermal challenge, which in the case of cetaceans would be continuous (Manger, 2006), the neurons and glia could be recruited to drive thermogenic processes, in addition to the functions normally associated with these cell types. This proposal is consistent with the known anatomical variances of the cetacean brain compared with other mammals (Kesarev, 1971; Manger, 2006; Patzke et al., 2015; Manger et al., 2010, 2012), the physiology and anatomy of cetacean sleep (Lyamin et al., 2008; Dell et al., 2016a,b), and the pragmatic view of cetacean behavior (Nikolskaya, 2005; Manger, 2013; Harley, 2013). Thus, we conclude that while the cetacean brain obviously provides adequate neural/cognitive processing to sustain life, it also exhibits the biological features that would allow it to produce sufficient heat to prevent suboptimal performance of the brain while under constant thermal challenge.

### Evolution of cetacean brains

The brains of cetaceans became both relatively and absolutely large around 20 million years after the ancestors of the modern cetacean fauna, the Archaeocetes, were already obligatory aquatic species. This enlargement in brain size occurred at the Archaeocete to modern cetacean fauna (the Neocete) faunal transition approximately 32 million years ago (mya) (Manger, 2006, 2013). Since this transition, the relative and absolute size of the brains of the Neocete have followed the same allometric scaling law of form (Manger, 2006). The thermogenetic hypothesis of cetacean brain evolution posits that, as this specific time point in the evolution of cetaceans (32 mya) coincides with significant drops in oceanic water temperatures as well as the loss of the warm, shallow, nutrient rich Tethys sea (Fordyce and Barnes, 1994; Whitemore, 1994; Zachos et al., 2001), the enlargement of the cetacean brain is an adaptive evolutionary response to thermal challenges (Manger, 2006, 2013). The data presented herein supports this notion and demonstrates that thermogenesis appears to be an augmented functional attribute of the cetacean brain in comparison to closely related artiodactyl mammals. Genetic studies on the cetacean nervous system demonstrate three major points of importance to our understanding of cetacean brain evolution (McGowen et al., 2012). First, 27 genes associated with the nervous system appear to have been positively selected for in the cetacean lineage including those specifically involved in sleep (McGowen et al., 2012), coinciding with the unusual sleep physiology of cetaceans that appears, in part, to be related to thermogenesis (Lyamin et al., 2008). Second, there appears to have been an ebb in the accumulation of genetic changes associated with the nervous system in the cetacean lineage (McGowen et al., 2012), which coincides with the stasis of the relative and absolute brain size of cetaceans following the Archaeocete – Neocete faunal transition (Manger, 2006, 2013). Third, seven mitochondrial expressed genes underwent positive selection in the cetacean lineage (McGowen et al., 2012), which coincides with the amplification and specialization of the expression of the UCP proteins 1, 4 and 5 shown herein. Thus, palaeoneurological, palaeoclimatological, genomic, neuroanatomical, neurochemical and neurophysiological studies of cetaceans all converge upon the concept that thermal pressures during the Archaeocete – Neocete faunal transition underlie the historical enlargement and current functionalities of the cetacean brain (Manger, 2006, 2013). This understanding of the evolution and functionality of the cetacean brain, which is reflected in their biogeographical distribution (Manger, 2006), may be of importance in providing a level of predictability to potential changes in the zoogeography of extant cetaceans in the face of rising ocean heat content associated with climate change (Cheng et al., 2019).

### Re-assessing large brain size evolution in mammals

The present study has broad reaching implications in terms of our understanding of the evolution of large brain size in mammals, including humans. By illustrating that it is possible to evolve a large brain for reasons not necessarily associated with a need for greater cognitive complexity, indicates that we should reassess our narratives regarding the evolution of large brains in humans, elephants and other mammals. In all of these situations, alternative explanations for increased brain size can be posited (Manger et al., 2013; González-Forero and Gardner, 2018). Most importantly, the current study emphasizes that, in terms of brain evolution and the resultant outcome, the starting point, this being what the brains of the ancestral species were like prior to enlargement, and any major environmental changes that occurred, are likely to be the best predictor of the functionality of the brain after enlargement. For cetaceans, the starting point was the Archaeocete brain, which, for animals that could grow to over 14 m in length, had a diminutive cerebral cortex with a total surface area of around 50 cm^2^ (Manger, 2006). On the other hand, the human brain evolved from an Australopithecine starting point, with brains quite similar to those seen in modern great apes, and thus the comparatively remarkable cognitive capacities of modern humans can be attributed in part to the enlargement of this ancestral brain, with the associated increases in neuronal complexity (Manger et al., 2013).

## Materials and Methods

### Specimens

We used brains obtained from three cetacean species (harbor porpoise – *Phocoena phocoena*, minke whale – *Balaenoptera acutorostrata*, and humpback whale – *Megaptera novaeangliae*) and 11 artiodactyl species (sand gazelle – *Gazella marica*, domestic pig – *Sus scrofa*, Nubian ibex – *Capra nubiana*, springbok – *Antidorcas marsupialis*, blesbok – *Damaliscus pygargus*, greater kudu – *Tragelaphus strepsiceros*, blue wildebeest – *Connochaetes taurinus*, dromedary camel – *Camelus dromedarius*, nyala – *Tragelaphus angasii*, river hippopotamus – *Hippopotamus amphibius*, and African buffalo – *Syncerus caffer*) (Table 1). All artiodactyl brains were perfusion fixed with 4% paraformaldehyde in 0.1 M phosphate buffer through the carotid arteries following euthanasia (Manger et al., 2009). The harbor porpoise specimens were perfusion fixed through the heart following euthanasia, while the minke whale and humpback whale brains were immersion fixed in 4% paraformaldehyde in 0.1 M phosphate buffer. All brains were then stored in an antifreeze solution at −20°C until use (Manger et al., 2009). All specimens were taken under appropriate governmental permissions, with ethical clearance provided by the University of the Witwatersrand Animal Ethics Committee (Clearance number 2008/36/1), which uses guidelines similar to those of the NIH regarding the use of animals in scientific research.

### Immunohistochemical staining

Blocks of tissue from the anterior cingulate (dorsal to the rostrum of the corpus callosum) and occipital cortex (presumably primary visual cortex) with underlying white matter were taken from each of the specimens. These were placed in a 30% sucrose in 0.1 M phosphate buffer solution at 4°C until equilibrated. The blocks were frozen in crushed dry ice, mounted on an aluminum stage and sectioned at 50 μm orthogonal to the pial surface. Alternate sections were stained for Nissl (with 1% cresyl violet), UCP1, UCP2, UCP3, UCP4, UCP5, dopamine-ß-hydroxylase (DBH) and tyrosine hydroxylase (TH). To investigate the presence of neural structures immunolocalizing uncoupling proteins, DBH and TH, we used standard immunohistochemical procedures with antibodies directed against UCP1, UCP2, UCP3, UCP4, UCP5, DBH and TH (see Supplementary Material for full immunohistochemical staining procedure). While immunolocalization for UCP1, UCP4, UCP5, DBH and TH were clear, only occasional cortical neurons were immunopositive for UCP2, and no immunolocalization could be detected for UCP3 in the species studied.

### Western immunoblotting

Protein expression for UCP1 and UCP4 was assayed using standard Western immunoblotting techniques. To verify the specificity of the UCP1 antibody for the UCP1 protein, we tested this antibody with rat brown fat. For the UCP4 antibody protein samples were extracted from the paraformaldehyde fixed tissue using the Qproteome FFPE Tissue Kit (Qiagen, Germany). The tissue blocks analyzed here were taken from the anterior cingulate and occipital cortex (as described above) and contained both gray and white matter. 30-40 mg of the sample were incubated in 100 μl of Extraction Buffer EXB Plus (Qiagen, Germany) containing 6% β-mercaptoethanol on ice for 5 min and mixed by vortexing. The samples were boiled for 20 min at 100°C and subsequently incubated at 50°C overnight with agitation at 300 rpm. The samples were then placed on ice for 1 min and centrifuged for 15 min at 14 000g at 4°C. The supernatant was transferred into clean tubes and the protein concentration was determined using the Bradford protein assay kit (Bio-Rad Laboratories, USA). The protein extracts (20 μg) were made soluble in sample buffer comprised of 0.0625 M Tris–HCl, pH 6.8, 10% glycerol, 2% SDS, 2.5% β-mercaptoethanol and 0.001% bromophenol blue, boiled at 95°C for 5 min and subjected to 12% SDS-polyacrylamide gel electrophoresis and transferred to polyvinylidene difluoride (PVDF) (Millipore) at 20 V/cm for 1h. Electrophoresis and protein transfer was achieved using Mini Trans-Blot Electrophoretic Transfer Cell (Bio-Rad Laboratories, Inc. USA). After the transfer the blots were blocked for 2 h in 1 x Animal-Free Blocker (SP-5030 Vector Labs, USA). The blots were incubated over night at 4°C under gentle agitation in the primary antibody solutions (1:300 goat anti-UCP1, Santa Cruz Biotechnology, sc-6528 or 1:300 goat anti-UCP4, Santa Cruz Biotechnology, sc-17582). The blots were washed for 3 x 10 min in 1 x Animal-Free Blocker and incubated for 1 h at room temperature in HRP-conjugated rabbit anti-goat secondary antibody (1:1000, Dako, USA) for 1 h. This was followed by 3 x 10 min washes with 50 mM Tris buffer, pH 7.2. The protein bands were detected using 3,3 ́-diaminobenzidine tetrahydrochloride hydrate (DAB) (Sigma, D5637). The blots were incubated in a solution containing 1mg/ml DAB in 50 mM Tris, pH 7.2 for 5 min at room temperature, followed by the addition of an equal amount of 0.02% hydrogen peroxide solution. Development was arrested by placing the blots in 50 mM Tris (pH 7.2) for 10 min, followed by two more 10 min rinses in distilled water.

### Stereological analysis

Using a design-based stereological approach we analyzed immunohistochemically stained sections in the grey matter of the anterior cingulate and occipital cortex, as well as the underlying white matter from these regions of 14 cetartiodactyl species. Regions of interest (ROI) were drawn from similar locations across species as supported by published anatomical descriptions of the cetacean and artiodactyl brain. Using a light microscope equipped with a motorized stage, digital camera, MicroBrightfield system (MBF Bioscience, USA) system and StereoInvestigator software (MBF Bioscience, version 2018.1.1; 64-bit), we quantified UCP1-immunoreactive neuron densities in the grey matter, UCP4-immunoreactive glia densities in the grey and white matter, and DBH- and TH-immunoreactive bouton densities in the grey and white matter of these cortical regions. Separate pilot studies for each immunohistochemical stain was conducted to optimise sampling parameters, such as the counting frame and sampling grid sizes, and achieve a coefficient of error (CE) below 0.1 (Dell et al., 2016a; Mouton, 2002; Gundersen, 1988; Gundersen and Jensen, 1987; West et al., 1991). In addition, we measured the tissue section thickness at every sampling site, and the vertical guard zone was determined according to tissue thickness to avoid errors/biases due to sectioning artefacts (Dell et al., 2016a; Mouton, 2002; Gundersen, 1988; Gundersen and Jensen, 1987; West et al., 1991). Supplementary tables S1-S4 provide details of the parameters used for each neuroanatomical region and stain and between the species in the current study. To estimate the ROI total number, we used the ‘Optical Fractionator’ probe.

UCP1- and UCP4-immunoreactive neuron and glia densities were obtained by sampling the cortical areas of interest and subjacent white matter with the aid of an optical disector. The cortex and white matter were outlined separately at low magnification (2X), and the optical disector was performed at 40X. UCP-immunoreactive neuron and glia density was calculated as the total number of UCP-immunoreactive neurons and glia divided by the product of surface area (x, y), the tissue sampling fraction, and the sectioned thickness (50 μm). The tissue sampling fraction was calculated as the ratio of the optical dissector height to mean measured section thickness. Given that overall cell density per unit volume is known to vary with differences in brain size, we calculated the percentage of UCP-immunoreactive neurons or glia, expressed as the ratio of UCP-immunoreactive neurons or glia to total neuronal or glial density for each region of interest, to standardize the data for cross species comparison. Using Nissl-stained sections we obtained estimates of neuronal and glial densities within the cortex and glial density within the white matter using optical disector probes combined with a fractionator sampling scheme (Mouton, 2002). A pilot study determined the optimal sampling parameters and grid dimensions to place dissector frames in a systematic-random manner. For DBH and TH bouton densities, ‘spot’ densities were calculated by multiplying the ROI area by the cut section thickness, and then using the generated volume as the denominator to the ROI estimated number. For all tissue sampled the optical fractionator was used while maintaining strict criteria, e.g. only complete boutons were counted, 63 X oil immersion, and obeying all commonly known stereological rules. The stereologic analyses presented here resulted in sampling an average of 118 counting frames per region of interest with a total of 13,053 counting frames investigated.

### Statistical analyses

We hypothesized that the percentage of cortical neurons immunoreactive to UCP1 were significantly different between artiodactyls and cetaceans. To test this hypothesis, we compared the proportion of UCP1 expression in the anterior cingulate and occipital cortex of 16 cetartiodactyls. For the anterior cingulate cortex, we sampled a total of 1 109 sampling sites (~ 100 sites per species) within the artiodactyl group and found that 36.83% of sampled cortical neurons were immunoreactive to UCP1. In comparison our cetacean sample consisted of 723 sampling sites (~ 145 sites per species), with 87.28% of the sampled cortical neurons immunoreactive to UCP1. For the occipital cortex, we sampled a total of 1 038 sites (~ 94 sites per species) within the artiodactyl group and found that 34% of sampled cortical neurons within the occipital cortex were immunoreactive to UCP1. The cetacean sample consisted of 723 sampling sites (~ 145 sites per species), and we found that 92.36% of the sampled cortical neurons were immunoreactive to UCP1.

To test if the respective underlying proportions were different between the sample groups, we conducted statistical hypothesis testing using the Two-Proportions Z-test as implemented in the R Programming language. Our Null hypothesis (*H*_o_) stated that there is no significant difference between the proportions of artiodactyl immunoreactive UCP1 sampled cortical neurons (π_1_) and the proportions of cetacean UCP1 sampled cortical neurons (π_2_) — that is, π_1_ – π_2_ = 0. The alternate hypothesis (*H*_1_) stated that there is a significant difference in these proportions such that π_1_ – π_2_ ≠ 0, with one of the proportions being either less than or greater than the other. We thus conducted a two-sided hypothesis test, with the significance level (α) set at 0.05 (i.e., *P*-values less than, or equal to, α, would reject the null hypothesis in favour of the alternate hypothesis). Based on these analyses the proportion of immunolabelled UCP1 cortical neurons were found to be significantly different between the groups, with cetaceans having a significantly higher proportion of UCP1-immunoreactive neurons in the anterior cingulate cortex (χ^2^ = 51.69; df =1, *P* = 6.49×10^−13^, 95% confidence interval = −0.122; −0.067) and occipital cortex (χ^2^ = 56.30; *P* = 6.21×10^−14^, 95% confidence interval = −0.114; −0.060).

We used a two sample T-test (as implemented in R) to test for significant differences in noradrenergic bouton density between cetaceans and artiodactyls. Cetaceans were found to have significantly higher mean DBH-immunoreactive bouton densities in the anterior cingulate cortex as compared to artiodactyls (*t* = −3.595; df =15, *P* = 0.011). Cetaceans were also found to have significantly higher mean DBH-immunoreactive bouton densities in the occipital cortex as compared to artiodactyls (*t* = − 4.546; df =15, *P* = 0.002). Similarly, we tested for significant differences in mean DBH bouton density in the underlying cortical white matter of cetaceans and artiodactyls. We did not find any significant differences in DBH-immunoreactive bouton density for the anterior cingulate (*t* =-0.597; df =15, *P* = 0.585) or occipital cortex (*t* = −0.08; df =15, *P* = 0.941).

To test for the effect of confounding variables on the significant differences observed in DBH bouton density in the cortex, we used an analysis of covariance controlling sequentially for the effect of cortical neuron density, cortical glia density and brain mass. Our analyses revealed that after adjusting for the density of cortical neurons cetaceans still had significantly higher DBH-immunoreactive bouton density in the anterior cingulate cortex (adjusted mean = 10.176) in comparison to artiodactyls (adjusted mean = 8.176) (*F* = 5.222; df =13, *P* = 0.041). Adjusting for the covariate cortical neuron density, resulted in a similar result for the occipital cortex (adjusted mean = 14.678) in comparison to artiodactyls (adjusted mean = 10.395) (*F* = 14.05; df =13, *P* = 0.00278). When controlling for the density of cortical glia, cetaceans also had significantly higher DBH-immunoreactive bouton densities in the anterior cingulate cortex (adjusted mean = 10.62) in comparison to artiodactyls (adjusted mean = 8.01) (*F* = 9.72; df =13, *P* = 0.00889). Similar results were found for the occipital cortex, with cetaceans having significantly higher DBH-immunoreactive bouton density (adjusted mean = 14.471) compared to artiodactyls (adjusted mean = 10.395) (*F* = 11.2; df =13, *P* = 0.00581). When controlling for brain mass, cetaceans were also found to have a significantly higher DBH-immunoreactive bouton densities in the anterior cingulate (adjusted mean = 11.36) in comparison to artiodactyls (adjusted mean = 7.75) (*F* = 11.06; df =13, *P* = 0.00604) as well as in the occipital cortex (cetacean adjusted mean = 15.406, artiodactyls adjusted mean = 10.055) (*F* = 11.85; df =13, *P* = 0.00488).

## Acknowledgments

This work was mainly supported by funding from the South African National Research Foundation (P.R.M.) and by a fellowship within the Postdoctoral-Program of the German Academic Exchange Service, DAAD (N.P.). The work was also supported by the Deanship of Scientific Research at the King Saud University through the EJ program (A.A.), the James S. McDonnell Foundation (P.R.H.), and the Swedish Research Council (04X-715, K.F.); P.R.M. and K.F. conceived the study and together with the remaining authors carried it out. N.P., A.B. and P.R.M. undertook the immunohistochemical staining, while N.P. did the western blotting. P.R.M. and K.Æ.K. collected the minke whale brains in Iceland, P.R.M. and A.B. collected the harbor porpoise brains in Greenland, P.R.M., A.N.A., N.C.B. and O.B.M. collected the artiodactyl brains from Saudi Arabia, P.R.M. and S.H.H. collected the artiodactyl brains from South Africa, P.R.M. and M.F.B. collected the hippopotamus brains from Denmark, P.R.H. provided the humpback whale brain tissue and M.A.S. and A.B. undertook the stereological and statistical analyses. P.R.M. prepared the paper, with all other authors making substantial intellectual input leading to the finished product; All data is available in the main text or the supplementary materials.

## Competing interests

Authors declare no competing interests;

## Supplementary Materials

### Immunohistochemical staining protocol

Blocks of tissue from the anterior cingulate (dorsal to the rostrum of the corpus callosum) and occipital cortex (presumably primary visual cortex) with underlying white matter were taken from each of the specimens. These were placed in a 30% sucrose in 0.1 M phosphate buffer solution at 4°C until equilibrated. The blocks were frozen in crushed dry ice, mounted on an aluminium stage and sectioned at 50 μm orthogonal to the pial surface. Alternate sections were stained for Nissl, UCP1, UCP2, UCP3, UCP4, UCP5, dopamine-ß-hydroxylase (DBH) and tyrosine hydroxylase (TH). To investigate the presence of neural structures immunolocalizing uncoupling proteins, DBH and TH, we used standard immunohistochemical procedures with antibodies directed against UCP1, UCP2, UCP3, UCP4, UCP5, DBH and TH. While immunolocalization for UCP1, UCP4, UCP5, DBH and TH were clear, only occasional cortical neurons were immunopositive for UCP2, and no immunolocalization could be detected for UCP3 in the species studied.

Sections used for the Nissl series were mounted on 0.5% gelatine-coated glass slides, cleared in a solution of 1:1 chloroform and absolute alcohol, then stained with 1% cresyl violet to reveal cell bodies. For the immunohistochemical staining, each section was treated with endogenous peroxidase inhibitor (49.2% methanol:49.2% 0.1 M PB: 1.6% of 30% H_2_O_2_) for 30 min and subsequently subjected to three 10 min 0.1 M PB rinses. Sections were then incubated for 2 h, at room temperature, in blocking buffer (containing 3% normal rabbit serum, NRS, for the UCP1-5 sections/3% normal horse serum, NHS, for the DBH sections/3% normal goat serum, NGS, for the TH sections, plus 2% bovine serum albumin and 0.25% Triton-X in 0.1 M PB). This was followed by three 10 min rinses in 0.1 M PB. The sections were then placed in the primary antibody solution that contained the appropriately diluted primary antibody in blocking buffer for 48 h at 4°C under gentle shacking. We used antibodies directed against UCP1 (Santa Cruz Biotechnology, C-17, sc-6528, Lot# D0411, goat polyclonal IgG, dilution 1:300), UCP2 (Santa Cruz Biotechnology, C-20, sc-6525, Lot# E0211, goat polyclonal IgG, dilution 1:300), UCP3 (Santa Cruz Biotechnology, C-20, sc-7756, Lot# A2511, goat polyclonal IgG, dilution 1:300), UCP4 (Santa Cruz Biotechnology, N-16, sc-17582, Lot# E2004, goat polyclonal IgG, dilution 1:300), UCP5 (Santa Cruz Biotechnology, Q-16, sc-50540, Lot# B1207, goat polyclonal IgG, dilution 1:300), DBH (Merck-Millipore, MAB308, mouse monoclonal IgG, dilution 1:4000) and TH (Merck-Millipore, AB151, rabbit polyclonal IgG, dilution 1:3000). This incubation was followed by three 10 min rinses in 0.1 M PB and the sections were then incubated in a secondary antibody solution (1:1000 dilution of biotinylated anti-goat IgG, BA-5000, Vector Labs, for UCP1-5 sections/1:1000 dilution of biotinylated anti-mouse IgG, BA 2001, Vector labs, for DBH sections/1:1000 dilution of biotinylated anti-rabbit IgG, BA-1000, Vector Labs, for TH sections, in a blocking buffer containing 3% NRS/NHS/NGS and 2% BSA in 0.1 M PB) for 2 h at room temperature. This was followed by three 10 min rinses in 0.1 M PB, after which sections were incubated for 1 h in avidin-biotin solution (at a dilution of 1:125, Vector Labs), followed by three 10 min rinses in 0.1 M PB. Sections were then placed in a solution of 0.05% 3,3’-diaminobenzidine (DAB) in 0.1 M PB for 5 min, followed by the addition of 3 ml of 3% hydrogen peroxide to each 1 ml of solution in which each section was immersed. Chromatic precipitation was visually monitored and verified under a low power stereomicroscope. Staining was allowed to continue until such time as the background stain was at a level that would assist architectural reconstruction and matching without obscuring the immunopositive neurons. Development was halted by placing the sections in 0.1 M PB, followed by two more rinses in 0.1M PB. To test for non-specific staining of the immunohistochemical protocol, in selected sections the primary antibody or the secondary antibody were omitted, which resulted in no staining of the tissue. The immunostained sections were then mounted on 0.5% gelatine coated glass slides, dried overnight, dehydrated in a graded series of alcohols, cleared in xylene and coverslipped with Depex. Digital photomicrographs were captured using Zeiss Axioshop and Axiovision software. No pixilation adjustments, or manipulation of the captured images were undertaken, except for the adjustment of contrast, brightness, and levels using Adobe Photoshop 7.

**Fig. S1.**
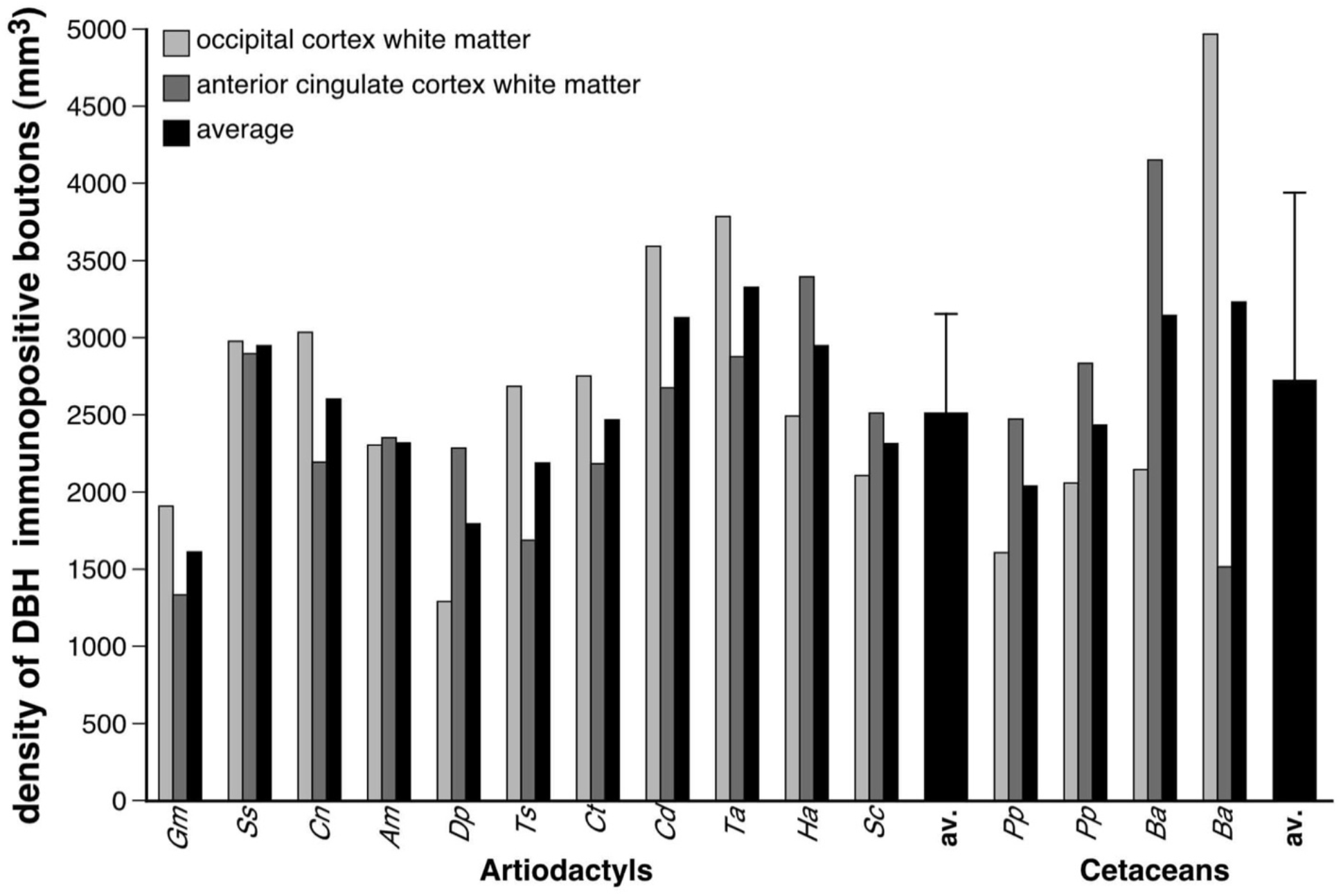
Quantification of noradrenergic bouton density in cetartiodactyl cortical white matter. The density of dopamine-ß-hydroxylase (DBH)-immunopositive boutons in the white matter below the anterior cingulate and occipital cortex was substantially lower than that observed in the corresponding grey matter (Tables 1, S3, error bars on average bars represent one standard deviation). No statistically significant differences were noted between artiodactyls and cetaceans. Depicted is a graphical representation of the results of the stereological analysis of the density of DBH-immunopositive boutons in the white matter of the occipital and anterior cingulate cortices of the species studied. ***Gm*** – sand gazelle, *Gazella marica*; ***Ss*** – domestic pig, *Sus scrofa*; ***Cn*** – Nubian ibex, *Capra nubiana*; ***Am*** – springbok, *Antidorcas marsupialis*; ***Dp*** – blesbok, *Damaliscus pygargus*; ***Ts*** – greater kudu, *Tragelaphus strepsiceros*; ***Ct*** – blue wildebeest, *Connochaetes taurinus*; ***Cd*** – dromedary camel, *Camelus dromedarius*; ***Ta*** – nyala, *Tragelaphus angasii*; ***Ha*** – river hippopotamus, *Hippopotamus amphibius*; ***Sc*** – African buffalo, *Syncerus caffer*; av. – average; ***Pp*** – harbor porpoise, *Phocoena phocoena*; ***Ba*** – minke whale, *Balaenoptera acutorostrata*.

**Fig. S2.**
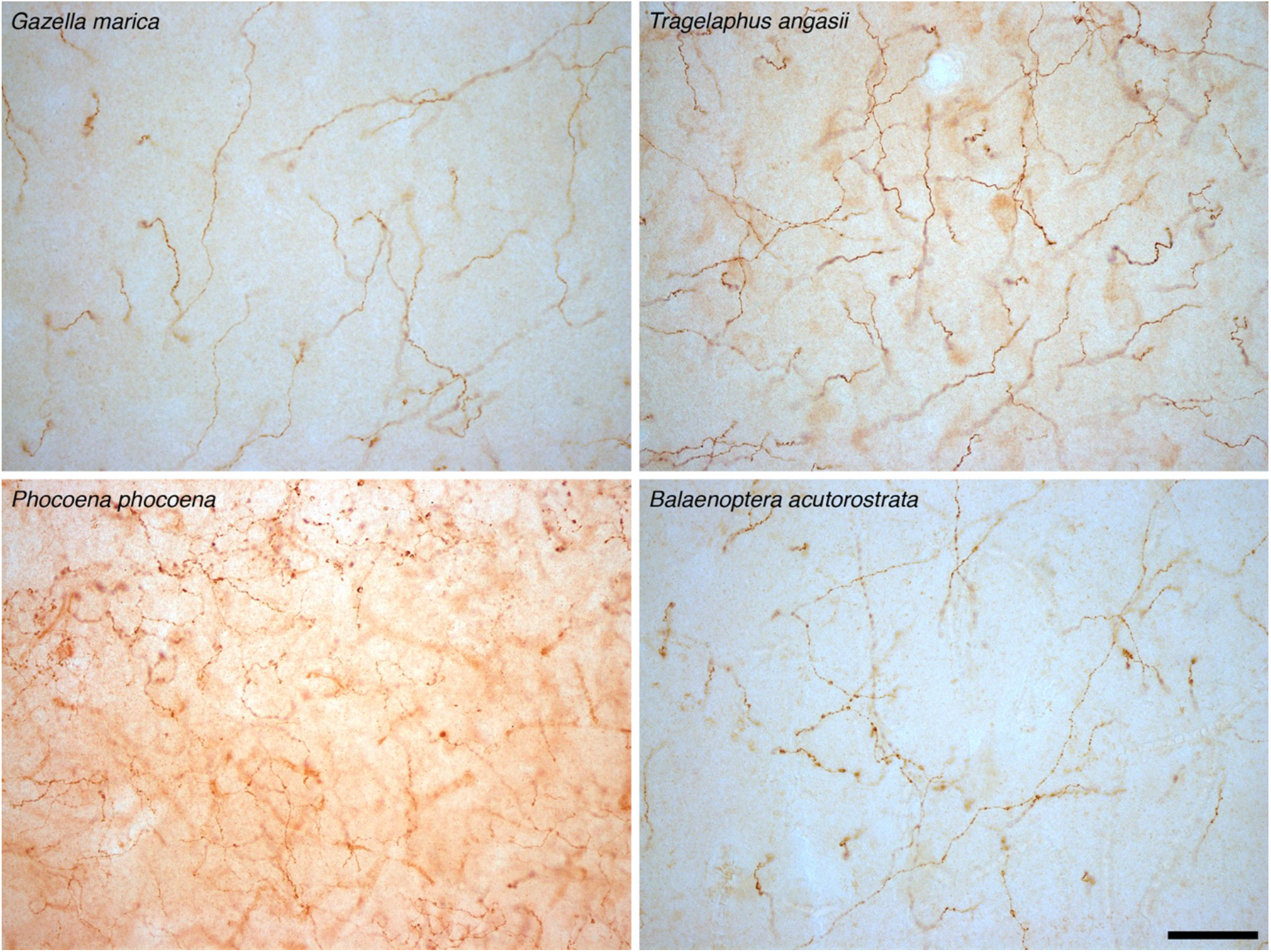
Tyrosine hydroxylase immunoreative boutons in cetartiodactyl cerebral cortex. In addition to revealing the density of catecholaminergic boutons in the cerebral cortex using dopamine-ß-hydroxylase (DBH), we stained these boutons with tyrosine hydroxylase (TH), marking an earlier stage in the catecholamine biosynthetic pathway. The average density of TH-immunoreactive boutons in the cortical grey matter of the artiodactyls studied was 7778 boutons/mm^3^ (range: 4073/mm^3^ in blesbok anterior cingulate cortex to 11704/mm^3^ in river hippopotamus occipital cortex). In cetacean cortical grey matter an average density of 14599 TH-immunoreactive boutons/mm^3^ was observed (range: 10717/mm^3^ in minke whale anterior cingulate cortex to 18247/mm^3^ in harbor porpoise occipital cortex) (Tables 1, S4). Using a two-sample *t*-test we compared TH-immunoreactive bouton density in the grey matter of the anterior cingulate and occipital cortex between artiodactyls and cetaceans. Cetaceans have significantly higher mean TH-immunoreactive bouton densities in both the anterior cingulate and occipital cortex compared to artiodactyls (anterior cingulate: *t* = −6.89; df =14, *P* = 0.00137; occipital cortex: *t* = −7.22; df =14, *P* = 0.0014). In the cortical white matter an average density of 1541 TH-immunoreactive boutons/mm^3^ was observed in artiodactyls, which was significantly (anterior cingulate: *t* = 0.53; df =14, *P* = 6.02 × 10-^1^; occipital: *t* =-4.09; df =14, *P* = 0.0016) lower than, the average TH-immunoreactive bouton density found in cetacean cortical white matter (1846 boutons/mm^3^) (Fig. S4). The photomicrographs presented here depict tyrosine hydroxylase (TH) immunostained axonal boutons in the cortical grey matter of *Gazella marica*, *Tragelaphus angasii*, *Phocoena phocoena*, and *Balaenoptera acutorostrata*. The scale bar = 50 μm and applies to all photomicrographs. Note the higher density of the TH-immunoreactive boutons in the cortical grey matter of cetaceans compared to the artiodactyls (see also Fig. S3).

**Fig. S3.**
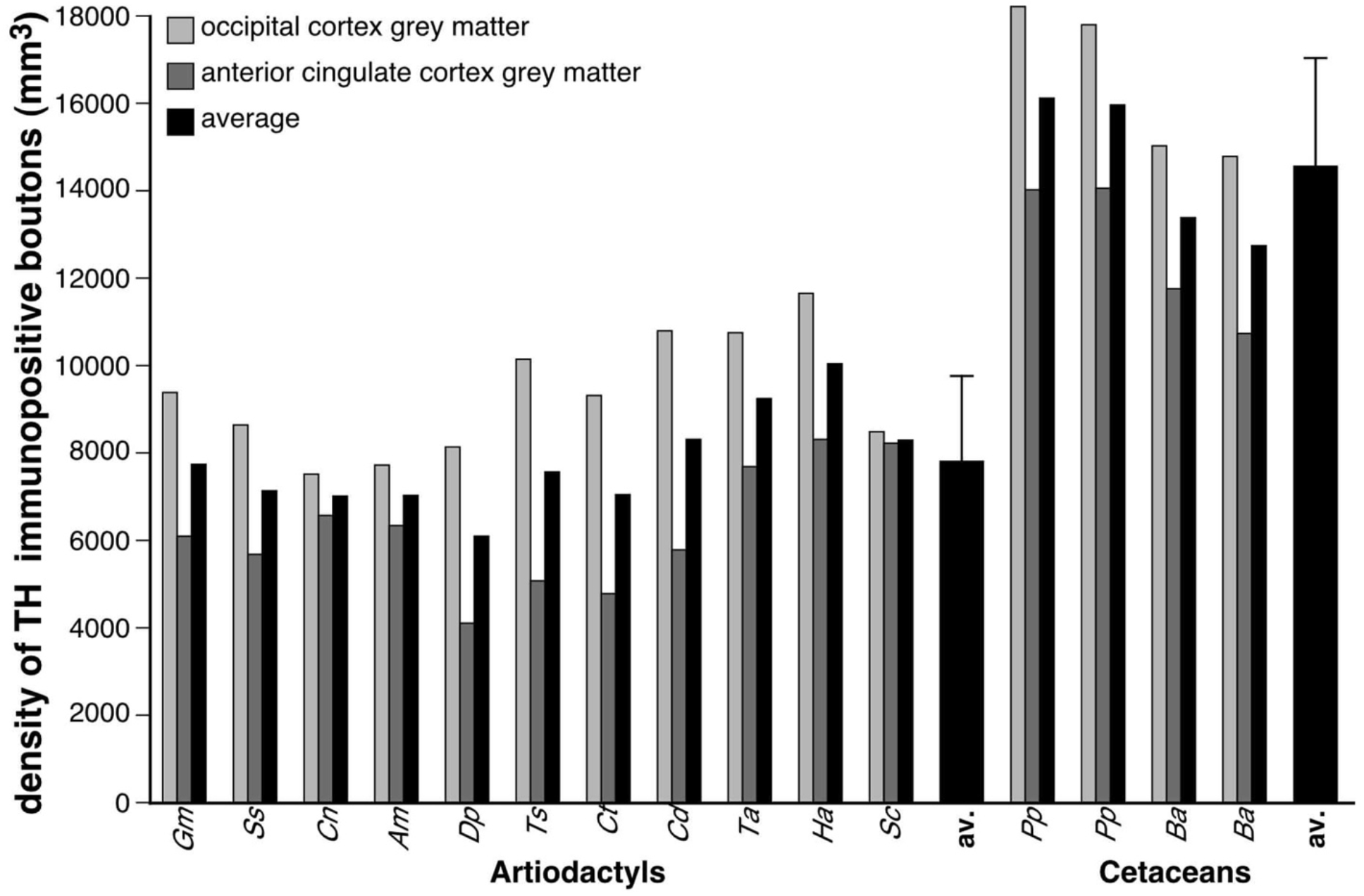
Quantification of tyrosine hydroxylase immunoreactive bouton density in cetartiodactyl cerebral cortex. Graphical representation of the results of the stereological analysis of the density of TH-immunopositive boutons in the grey matter of the occipital and anterior cingulate cortices of the species studied. Note that the density of these boutons is far higher in cetaceans than the artiodactyls (see legend of Fig. S2 for statistical results, error bars on average bars represent one standard deviation). ***Gm*** – sand gazelle, *Gazella marica*; ***Ss*** – domestic pig, *Sus scrofa*; ***Cn*** – Nubian ibex, *Capra nubiana*; ***Am*** – springbok, *Antidorcas marsupialis*; ***Dp*** – blesbok, *Damaliscus pygargus*; ***Ts*** – greater kudu, *Tragelaphus strepsiceros*; ***Ct*** – blue wildebeest, *Connochaetes taurinus*; ***Cd*** – dromedary camel, *Camelus dromedarius*; ***Ta*** – nyala, *Tragelaphus angasii*; ***Ha*** – river hippopotamus, *Hippopotamus amphibius*; ***Sc*** – African buffalo, *Syncerus caffer*; **av.** – average; ***Pp*** – harbor porpoise, *Phocoena phocoena*; ***Ba*** – minke whale, *Balaenoptera acutorostrata*.

**Fig. S4.**
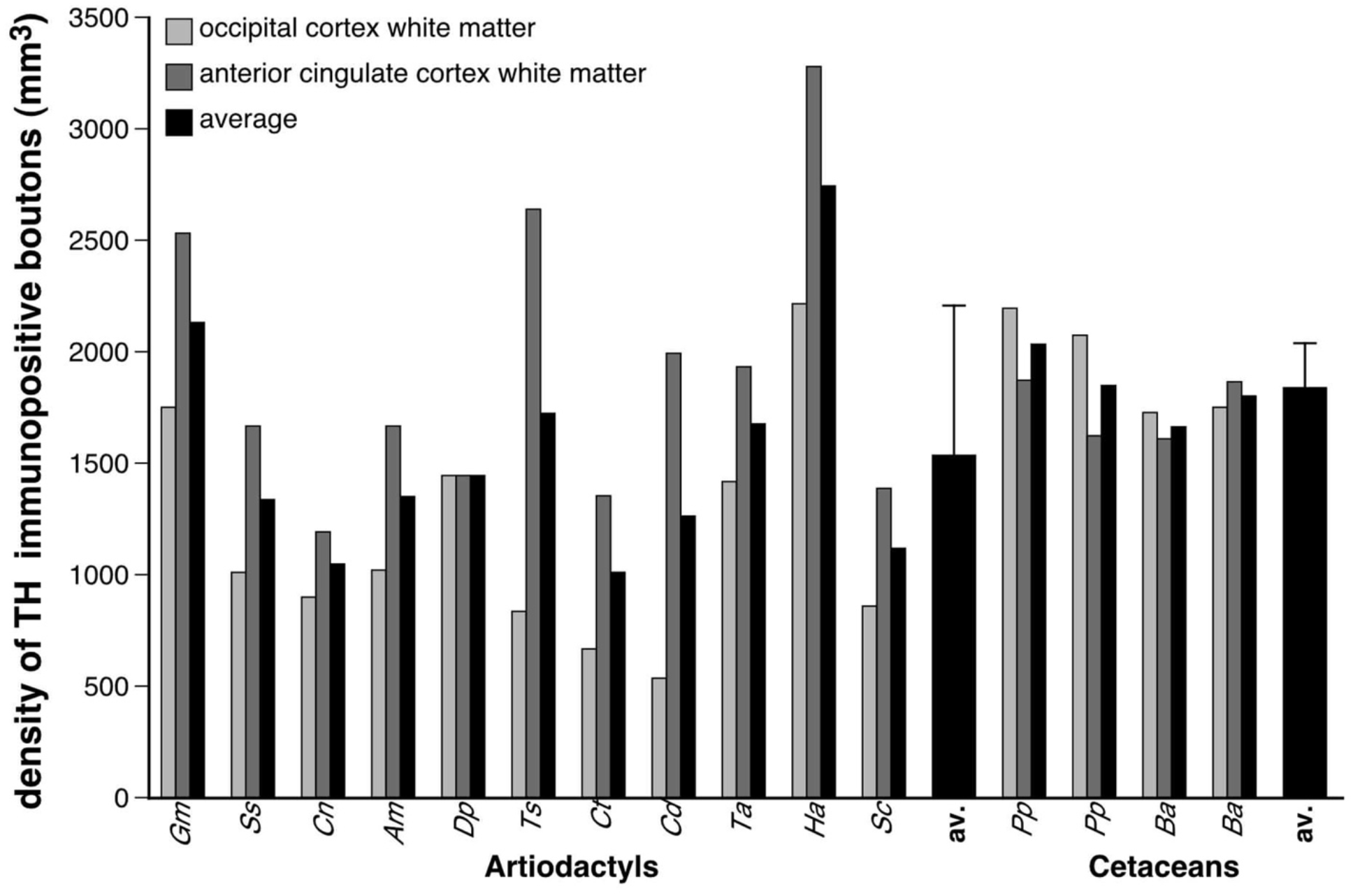
Quantification of tyrosine hydroxylase immunoreactive bouton density in cetartiodactyl subcortical white matter. Graphical representation of the results of the stereological analysis of the density of TH-immunopositive boutons in the white matter of the occipital and anterior cingulate cortices of the species studied. Note that the density of these boutons does not vary significantly across the species studied, although the average for cetaceans is slightly higher than that seen in the artiodactyls (see legend of Fig. S2 for statistical results, error bars on average bars represent one standard deviation). ***Gm*** – sand gazelle, *Gazella marica*; ***Ss*** – domestic pig, *Sus scrofa*; ***Cn*** – Nubian ibex, *Capra nubiana*; ***Am*** – springbok, *Antidorcas marsupialis*; ***Dp*** – blesbok, *Damaliscus pygargus*; ***Ts*** – greater kudu, *Tragelaphus strepsiceros*; *Ct* blue wildebeest, *Connochaetes taurinus*; ***Cd*** – dromedary camel, *Camelus dromedarius*; ***Ta*** – nyala, *Tragelaphus angasii*; ***Ha*** – river hippopotamus, *Hippopotamus amphibius*; ***Sc*** – African buffalo, *Syncerus caffer*; **av.** – average; ***Pp*** – harbor porpoise, *Phocoena phocoena*; ***Ba*** – minke whale, *Balaenoptera acutorostrata*.

**Fig. S5.**
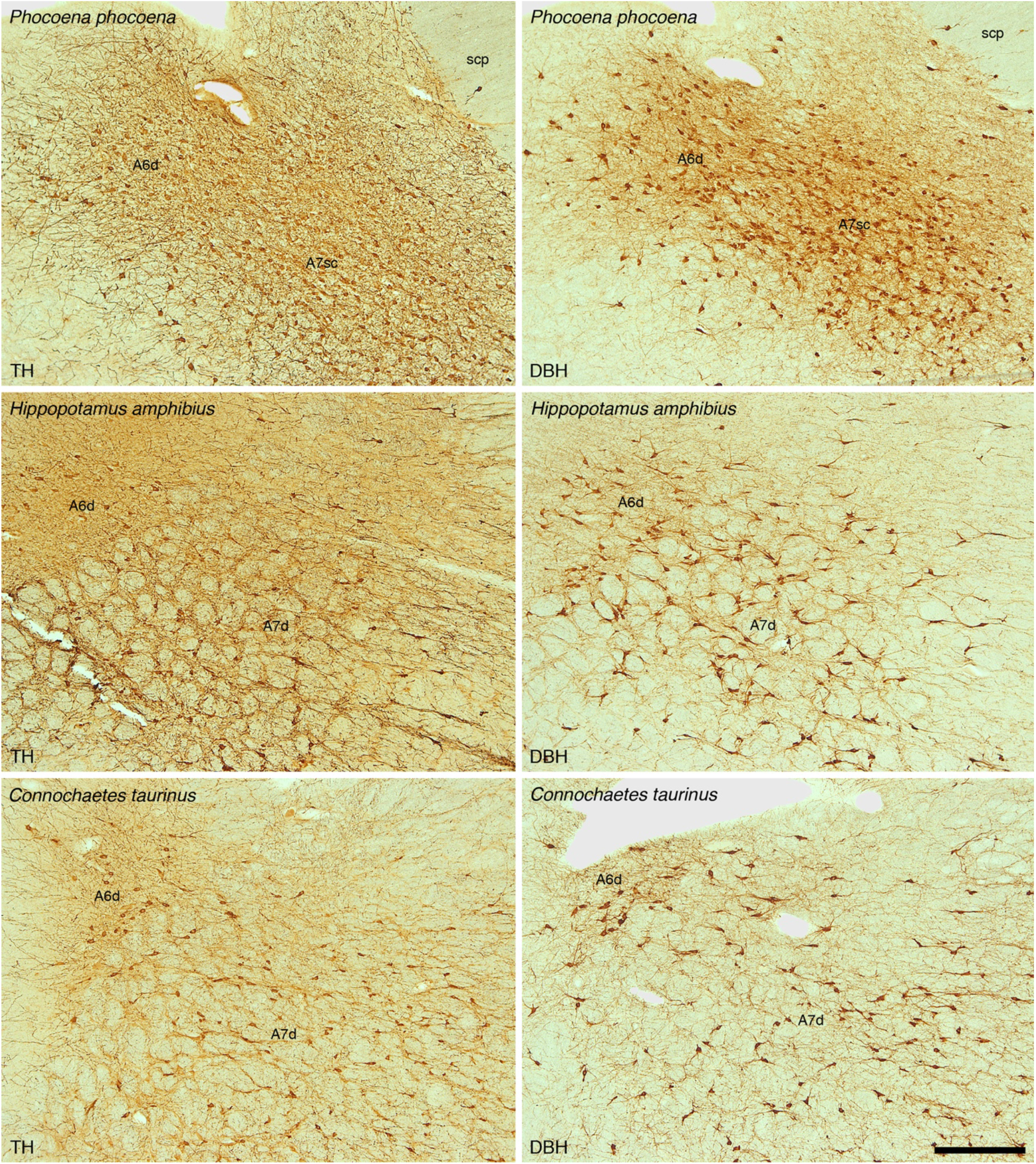
The locus coeruleus of cetartiodactyls. To support the concept that the noradrenergic innervation of the cerebral cortex arises from the locus coeruleus complex in the species studied, we examined the locus coeruleus with antibodies to tyrosine hydroxylase (TH) and dopamine-ß-hydroxylase (DBH). In all cases, the pattern of immunostaining indicates that the locus coeruleus of cetartiodactyls is the origin of noradrenergic projections throughout the brain. The photomicrographs provided here depict coronal sections through the locus coeruleus complex of the harbor porpoise (*Phocoena phocoena*), river hippopotamus (*Hippopotamus amphibius)* and blue wildebeest (*Connochaetes taurinus*) immunostained for TH (left column) and DBH (right column). Scale bar = 500 μm and applies to all. In all images dorsal is to the top and medial to the left. **A6d** – diffuse portion of locus coeruleus, **A7d** – diffuse portion of nucleus subcoeruleus, **A7sc** compact portion of nucleus subcoeruleus.

**Table S1.** Stereological parameters used in the estimation of UCP1-immunostained neuronal densities in the grey matter of the anterior cingulate (**AC**) and occipital (**OC**) cortices in the species studied.

| Species | Counting Frame Area ($\mu\text{m}^2$) | Sampling Grid Area ($\mu\text{m}^2$) | Disector height ($\mu\text{m}$) | Section cut thickness ($\mu\text{m}$) | Measured mounted thickness ($\mu\text{m}$) | Guard zone ($\mu\text{m}$) | Section interval | Number of sections | Number of sampling sites |
| --- | --- | --- | --- | --- | --- | --- | --- | --- | --- |
| Sand gazelle (AC) | 6400 | 1102500 | 4 | 50 | 13.4 | 2 | 1 | 3 | 40 |
| Sand gazelle (OC) | 6400 | 1102500 | 4 | 50 | 16.7 | 2 | 1 | 3 | 42 |
| Domestic pig (AC) | 6400 | 1102500 | 4 | 50 | 14.2 | 2 | 1 | 3 | 88 |
| Domestic pig (OC) | 6400 | 1102500 | 4 | 50 | 15.8 | 2 | 1 | 3 | 81 |
| Nubian ibex (AC) | 6400 | 1102500 | 4 | 50 | 14.6 | 2 | 1 | 3 | 109 |
| Nubian ibex (OC) | 6400 | 1102500 | 4 | 50 | 15.6 | 2 | 1 | 3 | 104 |
| Springbok (AC) | 6400 | 1102500 | 4 | 50 | 9.9 | 2 | 1 | 3 | 96 |
| Springbok (OC) | 6400 | 1102500 | 4 | 50 | 9.3 | 2 | 1 | 3 | 86 |
| Blesbok (AC) | 6400 | 1102500 | 4 | 50 | 11.4 | 2 | 1 | 3 | 139 |
| Blesbok (OC) | 6400 | 1102500 | 4 | 50 | 10.7 | 2 | 1 | 3 | 115 |
| Greater kudu (AC) | 6400 | 1102500 | 4 | 50 | 8.5 | 2 | 1 | 3 | 108 |
| Greater kudu (OC) | 6400 | 1102500 | 4 | 50 | 9.3 | 2 | 1 | 3 | 148 |
| Blue wildebeest (AC) | 6400 | 1102500 | 4 | 50 | 10.2 | 2 | 1 | 3 | 158 |
| Blue wildebeest (OC) | 6400 | 1102500 | 4 | 50 | 10 | 2 | 1 | 3 | 115 |
| Dromedary camel (AC) | 6400 | 1102500 | 4 | 50 | 16.8 | 2 | 1 | 3 | 54 |
| Dromedary camel (OC) | 6400 | 1102500 | 4 | 50 | 12.2 | 2 | 1 | 3 | 52 |
| Nyala (AC) | 6400 | 1102500 | 4 | 50 | 10.2 | 2 | 1 | 3 | 59 |
| Nyala (OC) | 6400 | 1102500 | 4 | 50 | 11.2 | 2 | 1 | 3 | 94 |
| River hippopotamus (AC) | 6400 | 1102500 | 4 | 50 | 8.6 | 2 | 1 | 3 | 129 |
| River hippopotamus (OC) | 6400 | 1102500 | 4 | 50 | 9.6 | 2 | 1 | 3 | 111 |
| African buffalo (AC) | 6400 | 1102500 | 4 | 50 | 12.9 | 2 | 1 | 3 | 108 |
| African buffalo (OC) | 6400 | 1102500 | 4 | 50 | 13.1 | 2 | 1 | 3 | 94 |
| Harbor porpoise 1 (AC) | 6400 | 1102500 | 4 | 50 | 10.6 | 2 | 1 | 3 | 100 |
| Harbor porpoise 1 (OC) | 6400 | 1102500 | 4 | 50 | 13.9 | 2 | 1 | 3 | 124 |
| Harbor porpoise 2 (AC) | 6400 | 1102500 | 4 | 50 | 11.2 | 2 | 1 | 3 | 112 |
| Harbor porpoise 2 (OC) | 6400 | 1102500 | 4 | 50 | 12.8 | 2 | 1 | 3 | 130 |
| Minke whale 1 (AC) | 6400 | 1102500 | 4 | 50 | 12.4 | 2 | 1 | 3 | 163 |
| Minke whale 1 (OC) | 6400 | 1102500 | 4 | 50 | 13.8 | 2 | 1 | 3 | 226 |
| Minke whale 2 (AC) | 6400 | 1102500 | 4 | 50 | 12.4 | 2 | 1 | 3 | 112 |
| Minke whale 2 (OC) | 6400 | 1102500 | 4 | 50 | 13.9 | 2 | 1 | 3 | 168 |

**Table S2.**
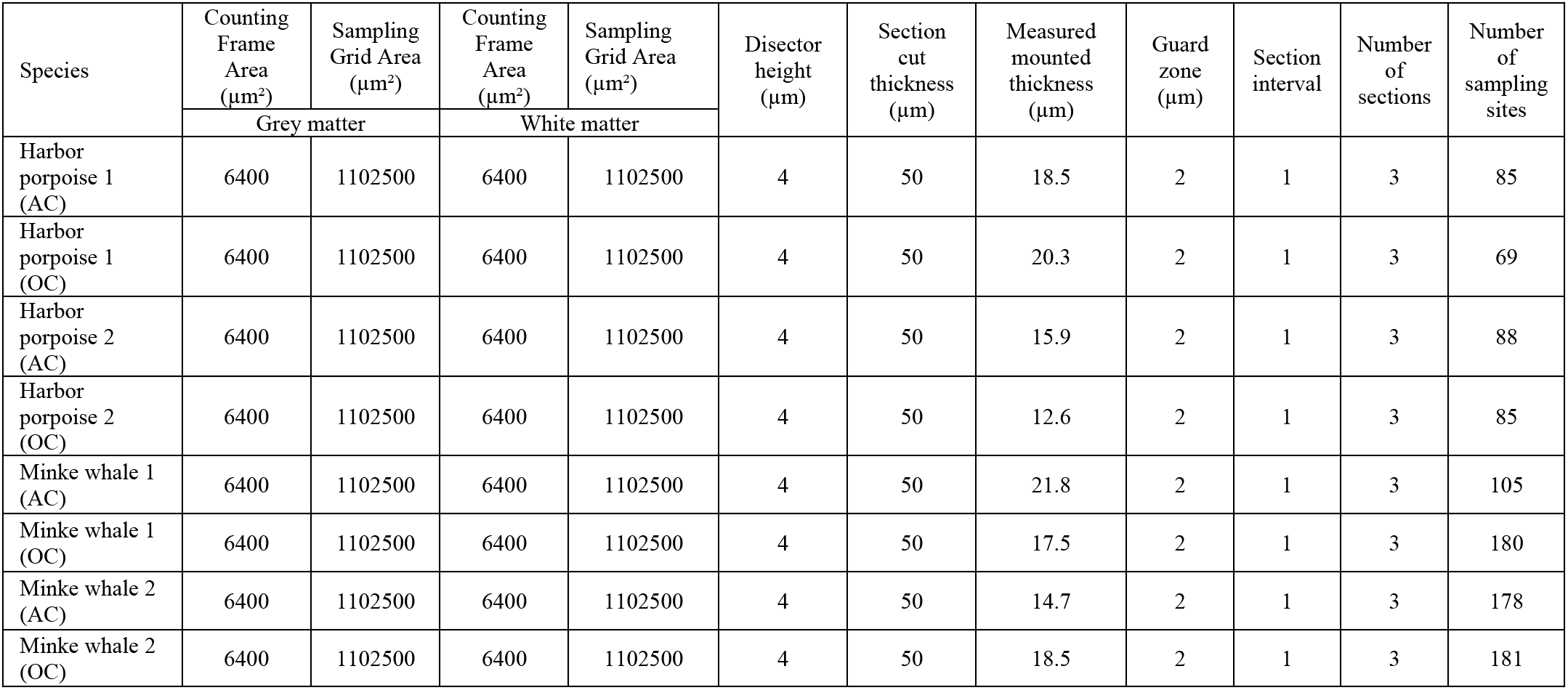
Stereological parameters used in the estimation of UCP4-immunostained glia densities in the grey and white matter of the anterior cingulate (**AC**) and occipital (**OC**) cortices in the cetacean species studied.

**Table S3.** Stereological parameters used in the estimation of dopamine-ß-hydroxylase (DBH)-immunoreactive bouton densities in the grey and white matter of the anterior cingulate (**AC**) and occipital (**OC**) cortices in the species studied.

| Species | Counting<br>frame size<br>(μm) | Sampling<br>grid size<br>(μm) | Counting<br>frame size<br>(μm) | Sampling<br>grid size<br>(μm) | Disector<br>height<br>(μm) | Section<br>cut<br>thickness<br>(μm) | Measured<br>mounted<br>thickness<br>(μm) | Guard<br>zone<br>(μm) | Section<br>interval | Number<br>of<br>sections | Number<br>of<br>sampling<br>sites |
| --- | --- | --- | --- | --- | --- | --- | --- | --- | --- | --- | --- |
|  | Grey matter |  | White matter |  |  |  |  |  |  |  |  |
| Sand gazelle<br>(AC) | 100 x 100 | 200 x 200 | 100 x 100 | 200 x 200 | 17 | 50 | 21.4 | 2 | 1 | 3 | 114 |
| Sand gazelle<br>(OC) | 100 x 100 | 200 x 200 | 100 x 100 | 200 x 200 | 17 | 50 | 22.3 | 2 | 1 | 3 | 112 |
| Domestic pig<br>(AC) | 100 x 100 | 200 x 200 | 100 x 100 | 200 x 200 | 16 | 50 | 20.5 | 2 | 1 | 3 | 118 |
| Domestic pig<br>(OC) | 100 x 100 | 200 x 200 | 100 x 100 | 200 x 200 | 16 | 50 | 20.3 | 2 | 1 | 3 | 116 |
| Nubian ibex<br>(AC) | 100 x 100 | 200 x 200 | 100 x 100 | 200 x 200 | 18 | 50 | 23.2 | 2 | 1 | 3 | 116 |
| Nubian ibex<br>(OC) | 100 x 100 | 200 x 200 | 100 x 100 | 200 x 200 | 18 | 50 | 22.4 | 2 | 1 | 3 | 119 |
| Springbok (AC) | 100 x 100 | 200 x 200 | 100 x 100 | 200 x 200 | 18 | 50 | 23.5 | 2 | 1 | 3 | 88 |
| Springbok (OC) | 100 x 100 | 200 x 200 | 100 x 100 | 200 x 200 | 18 | 50 | 22.7 | 2 | 1 | 3 | 91 |
| Blesbok (AC) | 100 x 100 | 200 x 200 | 100 x 100 | 200 x 200 | 19 | 50 | 23.7 | 2 | 1 | 3 | 107 |
| Blesbok (OC) | 100 x 100 | 200 x 200 | 100 x 100 | 200 x 200 | 19 | 50 | 24.0 | 2 | 1 | 3 | 93 |
| Greater kudu<br>(AC) | 100 x 100 | 250 x 250 | 100 x 100 | 250 x 250 | 17 | 50 | 22.1 | 2 | 1 | 3 | 116 |
| Greater kudu<br>(OC) | 100 x 100 | 250 x 250 | 100 x 100 | 250 x 250 | 17 | 50 | 21.8 | 2 | 1 | 3 | 107 |
| Blue wildebeest<br>(AC) | 100 x 100 | 250 x 250 | 100 x 100 | 250 x 250 | 20 | 50 | 24.5 | 2 | 1 | 3 | 114 |
| Blue wildebeest<br>(OC) | 100 x 100 | 250 x 250 | 100 x 100 | 250 x 250 | 20 | 50 | 24.6 | 2 | 1 | 3 | 101 |
| Dromedary<br>camel (AC) | 100 x 100 | 250 x 250 | 100 x 100 | 250 x 250 | 18 | 50 | 23.7 | 2 | 1 | 3 | 119 |
| Dromedary<br>camel (OC) | 100 x 100 | 250 x 250 | 100 x 100 | 250 x 250 | 18 | 50 | 22.8 | 2 | 1 | 3 | 107 |
| Nyala (AC) | 100 x 100 | 200 x 200 | 100 x 100 | 200 x 200 | 19 | 50 | 23.8 | 2 | 1 | 3 | 114 |
| Nyala (OC) | 100 x 100 | 200 x 200 | 100 x 100 | 200 x 200 | 19 | 50 | 24.3 | 2 | 1 | 3 | 107 |
| River<br>hippopotamus<br>(AC) | 100 x 100 | 250 x 250 | 100 x 100 | 250 x 250 | 18 | 50 | 23.1 | 2 | 1 | 3 | 96 |
| River<br>hippopotamus<br>(OC) | 100 x 100 | 250 x 250 | 100 x 100 | 250 x 250 | 18 | 50 | 22.4 | 2 | 1 | 3 | 102 |
| African buffalo<br>(AC) | 100 x 100 | 250 x 250 | 100 x 100 | 250 x 250 | 18 | 50 | 22.8 | 2 | 1 | 3 | 113 |
| African buffalo<br>(OC) | 100 x 100 | 250 x 250 | 100 x 100 | 250 x 250 | 18 | 50 | 22.5 | 2 | 1 | 3 | 84 |
| Harbor porpoise<br>1 (AC) | 100 x 100 | 250 x 250 | 100 x 100 | 250 x 250 | 19 | 50 | 23.7 | 2 | 1 | 3 | 113 |
| Harbor porpoise<br>1 (OC) | 100 x 100 | 250 x 250 | 100 x 100 | 250 x 250 | 19 | 50 | 23.2 | 2 | 1 | 3 | 99 |
| Harbor porpoise<br>2 (AC) | 100 x 100 | 250 x 250 | 100 x 100 | 250 x 250 | 16 | 50 | 20.4 | 2 | 1 | 3 | 97 |
| Harbor porpoise<br>2 (OC) | 100 x 100 | 250 x 250 | 100 x 100 | 250 x 250 | 16 | 50 | 20.5 | 2 | 1 | 3 | 113 |
| Minke whale 1<br>(AC) | 100 x 100 | 250 x 250 | 100 x 100 | 250 x 250 | 16 | 50 | 21.3 | 2 | 1 | 3 | 118 |
| Minke whale 1<br>(OC) | 100 x 100 | 250 x 250 | 100 x 100 | 250 x 250 | 16 | 50 | 20.8 | 2 | 1 | 3 | 121 |
| Minke whale 2<br>(AC) | 100 x 100 | 250 x 250 | 100 x 100 | 250 x 250 | 16 | 50 | 21.5 | 2 | 1 | 3 | 79 |
| Minke whale 2<br>(OC) | 100 x 100 | 250 x 250 | 100 x 100 | 250 x 250 | 16 | 50 | 20.6 | 2 | 1 | 3 | 121 |

**Table S4.** Stereological parameters used in the estimation of tyrosine hydroxylase (TH)-immunoreactive bouton densities in the grey and white matter of the anterior cingulate (**AC**) and occipital (**OC**) cortices in the species studied.

| Species | Counting<br>frame size<br>(µm) | Sampling<br>grid size<br>(µm) | Counting<br>frame size<br>(µm) | Sampling<br>grid size<br>(µm) | Disector<br>height<br>(µm) | Section<br>cut<br>thickness<br>(µm) | Measured<br>mounted<br>thickness<br>(µm) | Guard<br>zone<br>(µm) | Section<br>interval | Number<br>of<br>sections | Number<br>of<br>sampling<br>sites |
| --- | --- | --- | --- | --- | --- | --- | --- | --- | --- | --- | --- |
|  | Grey matter |  | White matter |  |  |  |  |  |  |  |  |
| Sand gazelle (AC) | 100 x 100 | 200 x 200 | 100 x 100 | 200 x 200 | 16 | 50 | 20.8 | 2 | 1 | 3 | 116 |
| Sand gazelle (OC) | 100 x 100 | 200 x 200 | 100 x 100 | 200 x 200 | 16 | 50 | 20.9 | 2 | 1 | 3 | 112 |
| Domestic pig (AC) | 100 x 100 | 200 x 200 | 100 x 100 | 200 x 200 | 16 | 50 | 21.2 | 2 | 1 | 3 | 113 |
| Domestic pig (OC) | 100 x 100 | 200 x 200 | 100 x 100 | 200 x 200 | 16 | 50 | 21.2 | 2 | 1 | 3 | 116 |
| Nubian ibex (AC) | 100 x 100 | 200 x 200 | 100 x 100 | 200 x 200 | 19 | 50 | 24 | 2 | 1 | 3 | 114 |
| Nubian ibex (OC) | 100 x 100 | 200 x 200 | 100 x 100 | 200 x 200 | 19 | 50 | 23.2 | 2 | 1 | 3 | 115 |
| Springbok (AC) | 100 x 100 | 200 x 200 | 100 x 100 | 200 x 200 | 16 | 50 | 21.8 | 2 | 1 | 3 | 89 |
| Springbok (OC) | 100 x 100 | 200 x 200 | 100 x 100 | 200 x 200 | 16 | 50 | 21.6 | 2 | 1 | 3 | 97 |
| Blesbok (AC) | 100 x 100 | 200 x 200 | 100 x 100 | 200 x 200 | 18 | 50 | 23.4 | 2 | 1 | 3 | 102 |
| Blesbok (OC) | 100 x 100 | 200 x 200 | 100 x 100 | 200 x 200 | 18 | 50 | 22.5 | 2 | 1 | 3 | 97 |
| Greater kudu (AC) | 100 x 100 | 250 x 250 | 100 x 100 | 250 x 250 | 17 | 50 | 21.8 | 2 | 1 | 3 | 117 |
| Greater kudu (OC) | 100 x 100 | 250 x 250 | 100 x 100 | 250 x 250 | 17 | 50 | 22.5 | 2 | 1 | 3 | 109 |
| Blue wildebeest (AC) | 100 x 100 | 250 x 250 | 100 x 100 | 250 x 250 | 17 | 50 | 21.9 | 2 | 1 | 3 | 114 |
| Blue wildebeest (OC) | 100 x 100 | 250 x 250 | 100 x 100 | 250 x 250 | 17 | 50 | 21.7 | 2 | 1 | 3 | 110 |
| Dromedary camel (AC) | 100 x 100 | 250 x 250 | 100 x 100 | 250 x 250 | 18 | 50 | 22.6 | 2 | 1 | 3 | 114 |
| Dromedary camel (OC) | 100 x 100 | 250 x 250 | 100 x 100 | 250 x 250 | 18 | 50 | 22.2 | 2 | 1 | 3 | 113 |
| Nyala (AC) | 100 x 100 | 200 x 200 | 100 x 100 | 200 x 200 | 17 | 50 | 22.4 | 2 | 1 | 3 | 116 |
| Nyala (OC) | 100 x 100 | 200 x 200 | 100 x 100 | 200 x 200 | 17 | 50 | 21.8 | 2 | 1 | 3 | 113 |
| River hippopotamus (AC) | 100 x 100 | 250 x 250 | 100 x 100 | 250 x 250 | 17 | 50 | 22.4 | 2 | 1 | 3 | 95 |
| River hippopotamus (OC) | 100 x 100 | 250 x 250 | 100 x 100 | 250 x 250 | 17 | 50 | 22.6 | 2 | 1 | 3 | 97 |
| African buffalo (AC) | 100 x 100 | 250 x 250 | 100 x 100 | 250 x 250 | 18 | 50 | 22.7 | 2 | 1 | 3 | 112 |
| African buffalo (OC) | 100 x 100 | 250 x 250 | 100 x 100 | 250 x 250 | 18 | 50 | 22.3 | 2 | 1 | 3 | 104 |
| Harbor porpoise 1 (AC) | 100 x 100 | 250 x 250 | 100 x 100 | 250 x 250 | 17 | 50 | 22.1 | 2 | 1 | 3 | 114 |
| Harbor porpoise 1 (OC) | 100 x 100 | 250 x 250 | 100 x 100 | 250 x 250 | 17 | 50 | 21.2 | 2 | 1 | 3 | 117 |
| Harbor porpoise 2 (AC) | 100 x 100 | 250 x 250 | 100 x 100 | 250 x 250 | 18 | 50 | 22.7 | 2 | 1 | 3 | 113 |
| Harbor porpoise 2 (OC) | 100 x 100 | 250 x 250 | 100 x 100 | 250 x 250 | 18 | 50 | 23.2 | 2 | 1 | 3 | 90 |
| Minke whale 1 (AC) | 100 x 100 | 250 x 250 | 100 x 100 | 250 x 250 | 18 | 50 | 23 | 2 | 1 | 3 | 118 |
| Minke whale 1 (OC) | 100 x 100 | 250 x 250 | 100 x 100 | 250 x 250 | 18 | 50 | 23.2 | 2 | 1 | 3 | 118 |
| Minke whale 2 (AC) | 100 x 100 | 250 x 250 | 100 x 100 | 250 x 250 | 18 | 50 | 23.3 | 2 | 1 | 3 | 118 |
| Minke whale 2 (OC) | 100 x 100 | 250 x 250 | 100 x 100 | 250 x 250 | 18 | 50 | 22.7 | 2 | 1 | 3 | 114 |

## Notes

### Competing Interest Statement

The authors have declared no competing interest.

